# Structure of the human inner kinetochore CCAN complex and its significance for human centromere organization

**DOI:** 10.1101/2022.01.06.475204

**Authors:** Marion E. Pesenti, Tobias Raisch, Duccio Conti, Ingrid Hoffmann, Dorothee Vogt, Daniel Prumbaum, Ingrid R. Vetter, Stefan Raunser, Andrea Musacchio

## Abstract

Centromeres are specialized chromosome loci that seed the kinetochore, a large protein complex that effects chromosome segregation. The organization of the interface between the kinetochore and the specialized centromeric chromatin, marked by the histone H3 variant CENP-A, remains incompletely understood. A 16-subunit complex, the constitutive centromere associated network (CCAN), bridges CENP-A to the spindle-binding moiety of the kinetochore. Here, we report a cryo-electron microscopy structure of human CCAN. We highlight unique features such as the pseudo GTPase CENP-M and report how a crucial CENP-C motif binds the CENP-LN complex. The CCAN structure has also important implications for the mechanism of specific recognition of the CENP-A nucleosome. A supported model depicts the interaction as fuzzy and identifies the disordered CCAN subunit CENP-C as only determinant of specificity. A more speculative model identifies both CENP-C and CENP-N as specificity determinants, but implies CENP-A may be in a hemisome rather than in a classical octamer.

## Introduction

The correct distribution of chromosomes from a mother cell to its daughters during mitosis and meiosis is of paramount importance for the stability of intra- and inter-generational genetic inheritance. A specialized proteinaceous complex named the kinetochore mediates the interaction of chromosomes and spindle microtubules and is essential for the success of this process. Kinetochores are complex macromolecular machines, consisting of approximately thirty core subunits, and are regulated at multiple levels to ensure errorless chromosome segregation (McKinley and Cheeseman, 2016; Musacchio and Desai, 2017).

Kinetochores assemble on a specialized chromosome segment known as the centromere. The histone H3 variant centromeric protein A (CENP-A) is considered the hallmark of centromeres (McKinley and Cheeseman, 2016; Mellone and Fachinetti, 2021; Talbert and Henikoff, 2020). It seeds kinetochores by recruiting the 16-subunit constitutive centromere associated network (CCAN) complex (Foltz et al., 2006; Izuta et al., 2006; Obuse et al., 2004; Okada et al., 2006) (Figure 1A). The CCAN is the heart of the inner (centromere-proximal) kinetochore and plays several crucial functions in kinetochore assembly and centromere maintenance. First, it provides several docking sites for the Knl1-Mis12-Ndc80 (KMN) network, a protein assembly of the outer (centromere-distal) kinetochore that mediates microtubules attachment and feedback control of the cell cycle (Musacchio and Desai, 2017). Second, it contributes to the inheritance of centromeres through cell division, which implies the continuous replenishment of CENP-A to compensate for its reduction during DNA replication (Jansen et al., 2007; Schuh et al., 2007). A basic organization comprising a CENP-A-based centromere and a CCAN-based inner kinetochore has undergone considerable evolutionary variation, but the general scheme remains recognizable in the vast majority of eukaryotes, including humans (Drinnenberg and Akiyoshi, 2017; Tromer et al., 2019; van Hooff et al., 2017).

**Figure 1.**
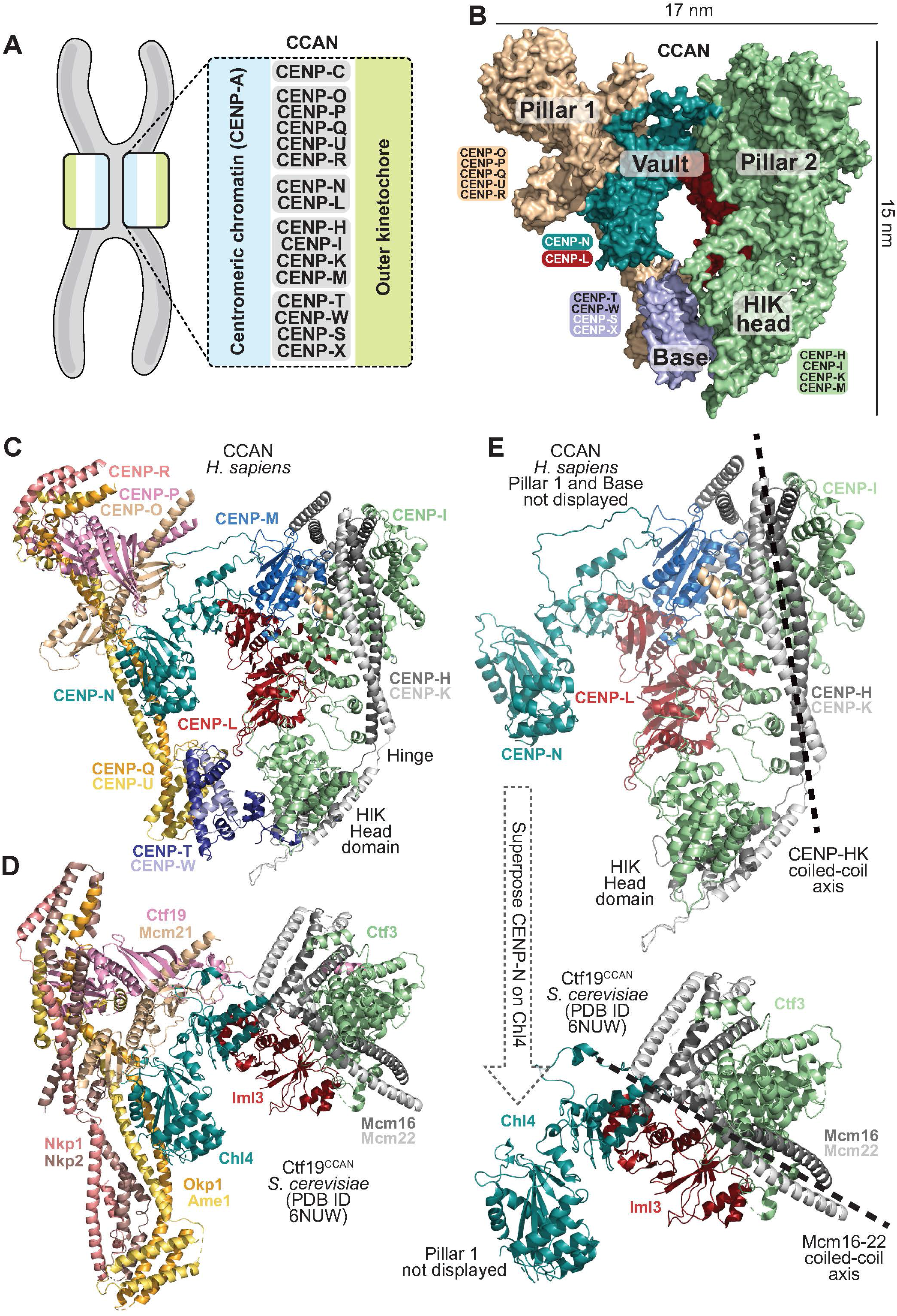
General organization of human CCAN. **A**) Scheme of kinetochore organization with presentation of CCAN subcomplexes. **B**) Surface model of CENP-16 complex coloured to identify distinct sub-modules. **C**) Cartoon model of human CCAN with all visible subunits. **D**) Cartoon model of the *S. cerevisiae* Ctf19^CCAN^ with same colouring scheme as for human subunits, as applicable (Hinshaw and Harrison, 2019; Yan et al., 2019). **E**) Cartoon models of human and yeast CCAN^Ctf19^ were superposed through CENP-N^Chl4^ and the resulting orientation of pillar 2 compared. This shows that pillar 2 adopts different orientations in the human and yeast complexes, more divergent in yeast, and more parallel to pillar 1 in humans. Only pillar 2 and the vault were displayed, while pillar 1 and the base were removed to enhance clarity.

In *Saccharomyces cerevisiae* and related yeasts, centromeres on different chromosomes are built on a conserved ≈125 base pairs (bps) segment of DNA. These centromeres are limited to a single specialized nucleosome defined by the presence of CENP-A (the *S. cerevisiae*’s CENP-A ortholog is known as Cse4) and are therefore defined as “point centromeres” (Bloom and Carbon, 1982; Fitzgerald-Hayes et al., 1982; Pluta et al., 1995). Most CCAN subunits are also identified in these organisms, where they are collectively referred to as the Ctf19 complex (henceforth Ctf19^CCAN^). Recent high-resolution cryo-electron microscopy structures of the *S. cerevisiae*’s Ctf19^CCAN^ complex revealed the reciprocal organization of subunits and a possible mode of interaction with an Cse4^CENP-A^ nucleosome (Hinshaw and Harrison, 2019, 2020; Yan et al., 2019; Zhang et al., 2020).

Contrary to the point centromere of *S. cerevisiae*, most eukaryotes have regional centromeres that extend over tens of thousands to even millions of DNA bases. These more complex centromeres often feature repetitive DNA sequences, such as the AT-rich, ∼171-base pairs (bps) *α*-satellite DNA repeats found at human kinetochores (McKinley and Cheeseman, 2016; Musacchio and Desai, 2017; Talbert and Henikoff, 2020). However, substantial evidence now supports the view that regional centromere assembly and inheritance is largely independent of DNA sequence. Rather, it is the presence of CENP-A and associated CCAN proteins that allows the propagation of centromeres through the cell-cycle-regulated recruitment of a specialized CENP-A loading machinery in a process that remains incompletely understood (Gambogi and Black, 2019; Mellone and Fachinetti, 2021).

The conservation of CCAN subunits in point and regional centromeres suggests that regional centromeres are modular, and assembled from the repetition of a basic “unit module”, similar to that existing in yeast, at multiple docking sites of the regional centromeres. Strong support for this idea has come from biochemical work on human proteins, which demonstrated the reconstitution of a discrete CCAN complex with purified components (McKinley et al., 2015; Pesenti et al., 2018; Walstein et al., 2021; Weir et al., 2016). Furthermore, a low-resolution structural by negative-stain electron microscopy provided a first view of the human CCAN complex, revealing a structure that in its outline appears similar to that in *S. cerevisiae* and related yeasts (Hamilton et al., 2019; Kixmoeller et al., 2020; Pesenti et al., 2018).

The detailed organization of centromeric chromatin has been debated, with models ranging from a right-handed hemisome or a hexasome for the *S. cerevisiae* centromere to classical left-handed octameric CENP-A containing nucleosome in humans (Black and Cleveland, 2011; Talbert and Henikoff, 2020). Octameric CENP-A mono-nucleosomes assemble *in vitro* and have been extracted from nuclease-treated chromatin (Hasson et al., 2013; Nechemia-Arbely et al., 2017; Tachiwana et al., 2011). Previous studies, however, have shown that native centromeres may disassemble when chromatin is trimmed to mono-nucleosomes (Ando et al., 2002). Thus, while there is substantial evidence for an octameric left-handed CENP-A nucleosome, it is important to assess the organization of centromeric chromatin in complexes that retain fundamental properties of the native structure. Proteins mediating specific CENP-A recognition by CCAN have a crucial role in this regard. Two CCAN proteins have emerged for specific recognition of CENP-A: CENP-C and CENP-N. CENP-C binds CENP-A nucleosomes through two related motifs, the central region and the CENP-C motif. CENP-N, on the other hand, recognizes the L1 loop of CENP-A, where the sequence of CENP-A diverges from the sequence of H3 (Ali-Ahmad et al., 2019; Allu et al., 2019; Ariyoshi et al., 2021; Carroll et al., 2010; Carroll et al., 2009; Chittori et al., 2018; Guo et al., 2017; Kato et al., 2013; Pentakota et al., 2017; Tian et al., 2018; Walstein et al., 2021)

Here, we report cryo-electron microscopy (cryo-EM) structures of human CCAN assemblies comprising 16 subunits, designated CENP-16, and including the N-terminal region of CENP-C^1-544^ and the subcomplexes CENP-O/CENP-P/CENP-Q/CENP-U/CENP-R (CENP-OPQUR complex), CENP-N/CENP-L (CENP-LN), CENP-H/CENP-I^57-765^/CENP-K/CENP-M (CENP-HIKM), and CENP-T/CENP-W/CENP-S/CENP-X (CENP-TWSX). We show that the human CCAN structure is similar to the yeast Ctf19^CCAN^ structure in its outline, but diverges from it in crucial aspects that have important implications for nucleosome binding. Furthermore, we report that previous structures of the N-terminal region of CENP-N in complex with the octameric CENP-A nucleosome (Allu et al., 2019; Chittori et al., 2018; Pentakota et al., 2017; Tian et al., 2018) are incompatible with the structure of human CCAN. We discuss various models of centromere organization that might reconcile these observations and their implications for centromere biology.

## Results

### General organization of CCAN

We built on our previous biochemical reconstitution of the human kinetochore (Pesenti et al., 2018; Walstein et al., 2021; Weir et al., 2016) to generate CENP-16 from stable individual subcomplexes in preparation for high-resolution cryo-EM data collection (Figure 1 – Supplement 1 and Figure 1 – Supplement 2). Two cryo-EM datasets, including one of pure CENP-16 (dataset I) and one of CENP-14 (lacking CENP-SX) with DNA and CENP-A:H4 (dataset II), were processed independently as outlined in Figure 1 – Supplement 3 and Figure 1 – Supplement 4 (see also Table S1 and Methods). We obtained reconstructions for both datasets. However, only the cryo-EM structure derived from dataset I yielded a resolution range (3.7 Å in the centre with lower resolution in peripheral parts) that allowed reliable model building (Figure 1 – Supplement 3-5).

Molecular models of CCAN subunits were either available from previous structural work (CENP-M, CENP-N), or were generated by homology modelling based on structures of yeast CCAN (Hinshaw and Harrison, 2019, 2020; Yan et al., 2019; Zhang et al., 2020), and in later phases also based on AlphaFold2 in the variants Colabfold and AF2-Multimer (Evans et al., 2021; Jumper et al., 2021; Mirdita et al., 2021). Models were fitted in the density using a combination of manual and automated fitting methods (see Methods). The final high-resolution model of CENP-16 obtained with dataset I encompasses 14 of the 16 subunits. CENP-S and CENP-X (CENP-SX), which require CENP-TW for incorporation into CENP-16 (Figure 1 – Supplement 1E-F), could not be modelled as we could not identify a density for these subunits, suggesting they are disordered or absent from the particles). The final model has a molecular mass of ≈450 kDa and consists of ≈25000 atoms.

The organization of CENP-16 can be rationalized as consisting of two “pillars” connected by a “vault” and a “base” (Figure 1B). Pillars 1 and 2 consist of CENP-OPQUR and CENP-HIKM, respectively. The vault consists of CENP-LN. The base consists of CENP-TW, whose position is more clearly defined in the lower resolution map from dataset II. This may indicate an effect from the presence of DNA and CENP-A:H4 in this sample, but neither was recognizable in density maps. A comparison of human CCAN with the yeast Ctf19^CCAN^ reveals overall similarities (Figure 1C-D), further reinforcing the concept that chromatin of regional centromeres consists of a repetitive CCAN unit module resembling the single Ctf19^CCAN^ of the yeast point centromere. Nonetheless, there are also crucial differences with important implications for our understanding of centromeric chromatin, as described below. Most notably, the angle between pillar 1 and pillar 2 is much wider in the yeast structure than in the human structure, as shown by taking as reference the axis of the CENP-HK coiled-coil (Figure 1E). As discussed below, this difference results from several factors, including the presence of CENP-M and a more closed CENP-LN vault in the human complex relative to the yeast complex (see also Figure 6 below).

The resolution of our maps is highest in the central part of the complex where the vault interacts with pillars 1 and 2, and decreases in more peripheral regions (Figure 1 – Supplement 4). Nevertheless, there is significant density, especially in maps obtained with dataset II, for the complex of the CENP-I N-terminal domain with the CENP-HK C-terminal domain, which together form a “HIK head domain” connected to the rest of pillar 2 by a short hinge (Figure 1C-D). The HIK head domain, which in the yeast complex is visible in two different conformations in the structures of Ctf19^CCAN^ dimers and in the structure of the Ctf19^CCAN^ complex with the Cse4^CENP-A^ nucleosome, associates with the histone fold domains (HFDs) of CENP-T and CENP-W (the “base”). These were modelled with AF2 and also with reference to recent crystal structures of the Cnn1^CENP-T^/Wap1^CENP-W^ complex bound to the *S. cerevisiae* HIK head (PDB ID 6WUC and 6YPC) (Hinshaw and Harrison, 2020; Zhang et al., 2020) (Figure 1C). A structure-based alignment of yeast and human CCAN^Ctf19^ subunits is supplied in Figure 1 – Supplement 6.

### CENP-R, CENP-C, and CENP-M

CENP-R caps the “northern” globular head of pillar 1 (Figure 1C and Figure 2A). Density for CENP-R is limited to a region comprising residues Phe86-Gln149 as assigned by the AF2 model, consisting of two α-helices and a short helical connector (Figure 2B). The rest of the structure is predicted as intrinsically disordered. The CENP-R helices pack closely against a short pair of helices at the C-terminus of the CENP-QU sub-complex. Overall, their position is roughly equivalent to that of α-helices of Nkp1 and Nkp2 in the related structure of the *S. cerevisiae* Ctf19^CCAN^ complex (Hinshaw and Harrison, 2019; Yan et al., 2019) (Figure 1D), possibly suggesting a distant evolutionary relationship. However, Nkp1 and Nkp2 accompany the entire length of pillar 1 in Ctf19^CCAN^, tightly interacting with Okp1^CENP-U^ and Ame1^CENP-Q^, whereas the interaction of CENP-R is limited to the head of pillar 1. As a result, pillar 1 is thinner in human CCAN (Figure 1C-D).

**Figure 2.**
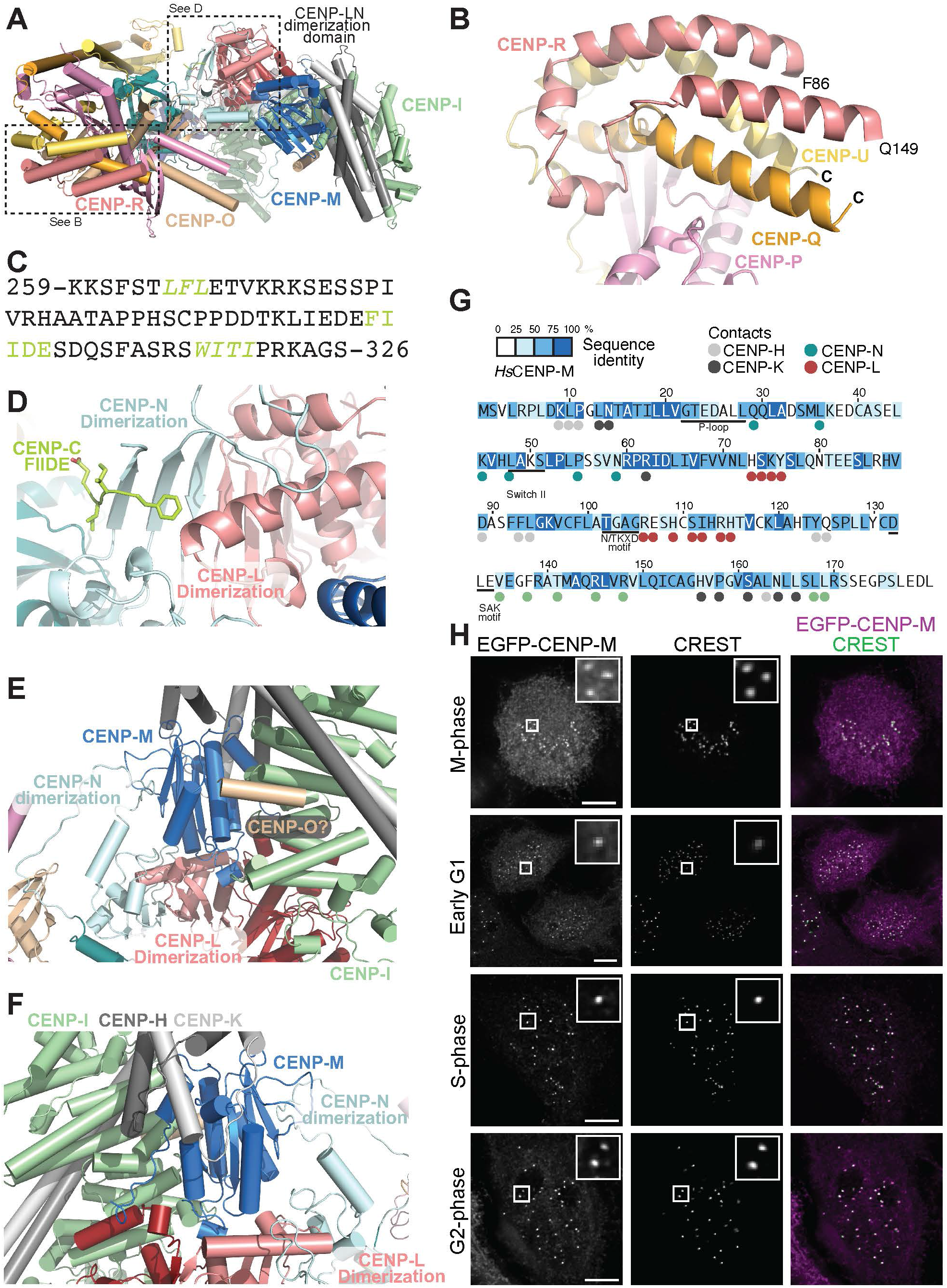
CENP-C, CENP-M, and CENP-R. **A**) Cartoon showing an overall view of human CCAN from above, and displaying the “knob” domain of pillar 1, the dimerization domain of the vault subunits CENP-LN, and the upper domain of pillar 2, where CENP-M resides. **B**) Close-up view of the two helices of CENP-R with initial and final visible residues, and the connecting helical segment. **C**) Sequence of HsCENP-C within the CCAN binding region. Two sequences shown in green and italics identify motifs previously shown to interact with CENP-NL and CENP-HIKM (Klare et al., 2015). The FIIDE sequence interacts with the CENP-LN dimerization domain. **D**) Cartoon model of the CENP-LN dimerization domain with the bound FIIDE sequence of CENP-C shown in sticks. **E**) Embedding of CENP-M in a network of interactions between pillar 2 and the vault. **F**) A rotated view showing additional CENP-M interactions. **G**) Sequence of CENP-M with displayed conservation from 12 distant CENP-M orthologs and contacts with neighbouring subunits. The figure was adapted from reference (Basilico et al., 2014). CENP-M residues in contact with other CCAN subunits are indicated with the same colour used for representation of the subunit. **H**) Localization of an EGFP-CENP-M construct in HeLa cells during the indicated phases of the cell cycle demonstrates continuity of localization.

Most of the 943-residue protein CENP-C is predicted as intrinsically disordered. Previous work rationalized CENP-C as a blueprint for kinetochore assembly, as the protein contains, from the N- to the C-terminus, a succession of linear binding motifs for other kinetochore proteins ordered along the outer to inner kinetochore axis (Klare et al., 2015; Walstein et al., 2021). The motifs begin near the N-terminus with an interaction motif for the outer kinetochore MIS12 complex (Gascoigne et al., 2011; Screpanti et al., 2011), followed by binding motifs for the CCAN subunits CENP-LN and CENP-HIKM (comprised between residues 259 and 326 and highlighted in green in Figure 2C) (Klare et al., 2015; Pentakota et al., 2017), and further down for the CENP-A nucleosome, with two related CENP-A binding motifs in humans, the central region (residues 515-535) and the CENP-C motif (735-755) (Ariyoshi et al., 2021; Guo et al., 2017; Kato et al., 2013; Walstein et al., 2021). Finally, CENP-C dimerizes through its only sizable folded region, the C-terminal cupin domain (Chik et al., 2019; Cohen et al., 2008; Medina-Pritchard et al., 2020; Walstein et al., 2021).

CENP-16 incorporates CENP-C residues 1-544 (CENP-C^1-544^), but there is no discernible CENP-C density except for a Phe-Ile-Ile-Asp-Glu (303-FIIDE-307) fragment. In previous work, we found biochemically that this motif is crucial for CENP-LN recruitment to human kinetochores and that it may bind near the CENP-LN dimerization domain (Pentakota et al., 2017). This prediction is fully confirmed by the structural analysis (Figure 2D). The sequence assignment of CENP-C into this short stretch of density was based on a highly significant AF2 prediction for this interaction. Supported by AF2 predictions, we also tentatively assigned a predicted single α-helix at the N-terminus of CENP-O to an unaccounted density at the interface of CENP-HK, CENP-I, and CENP-M (Figure 2E). The region containing the 265-LFL-268 motif of CENP-C found to interact with CENP-HIKM (Klare et al., 2015) is also predicted as α-helix by AF2 and could in principle also fit in the unaccounted helical density, but AF2 did not predict any significant interactions between this CENP-C region and the CENP-HIKM subcomplex.

The pseudo GTPase CENP-M, which is unable to bind and hydrolyse GTP (Basilico et al., 2014), binds near the CENP-LN dimerization domains at the vault’s apex (Figure 1C). Using conserved interfaces, CENP-M also wedges against CENP-I and CENP-HK, generating a very robust network of interactions that bury collectively more that 3300 Å^2^ and greatly stabilizes CENP-16 (Figure 2E-G and Figure 2 – Supplement 1A). As CENP-M resides at the kinetochore throughout the cell cycle (Figure 2H and Figure 2 – Supplement 2), we suspect its stabilizing function is constitutive and unregulated. Thus, collectively, pillar 2 is much more deeply connected to the CENP-LN vault of CCAN in the human complex than in the Ctf19^CCAN^ complex, where the trimer of Ctf3^CENP-I^, Mcm16^CENP-H^, and Mcm22^CENP-K^ connects to the rest of Ctf19^CCAN^ exclusively through a small interface (∼550 Å^2^) between Ctf3^CENP-I^ and Iml3^CENP-L^ (Figure 2 – Supplement 1B).

### CENP-16 binds DNA

A surface representation of CENP-16 demonstrates the presence of a central tunnel surrounded by the CENP-TW base and the CENP-LN vault domains, and further extended by pillar 2 on the front and pillar 1 on the back of the structure (Figure 3A). Several considerations, some of which discussed more extensively below, converge on identifying the central tunnel of CENP-16 as a DNA-binding interface. Lined with several positively charged residues from CENP-L and CENP-N, and with an internal diameter of ≈27 Å, the vault seems ideally suited to surround the negatively charged backbone of a double-stranded (ds) DNA filament with a diameter of ≈20 Å. CENP-TW in the CCAN base also expose the DNA-binding interface of the histone fold domains (HFDs), which flank positive patches on CENP-I in the HIK head domain in the front (Figure 3B, *left*) and on CENP-QU in the back (Figure 3B, *right*). Another positive patch on CENP-N, contributed by residues in the α6 helix, including K102, K110, and R114 (Figure 3B, middle) has been recently implicated in CENP-A nucleosome stacking by the CENP-N N-terminal domain (Zhou et al., 2021).

**Figure 3.**
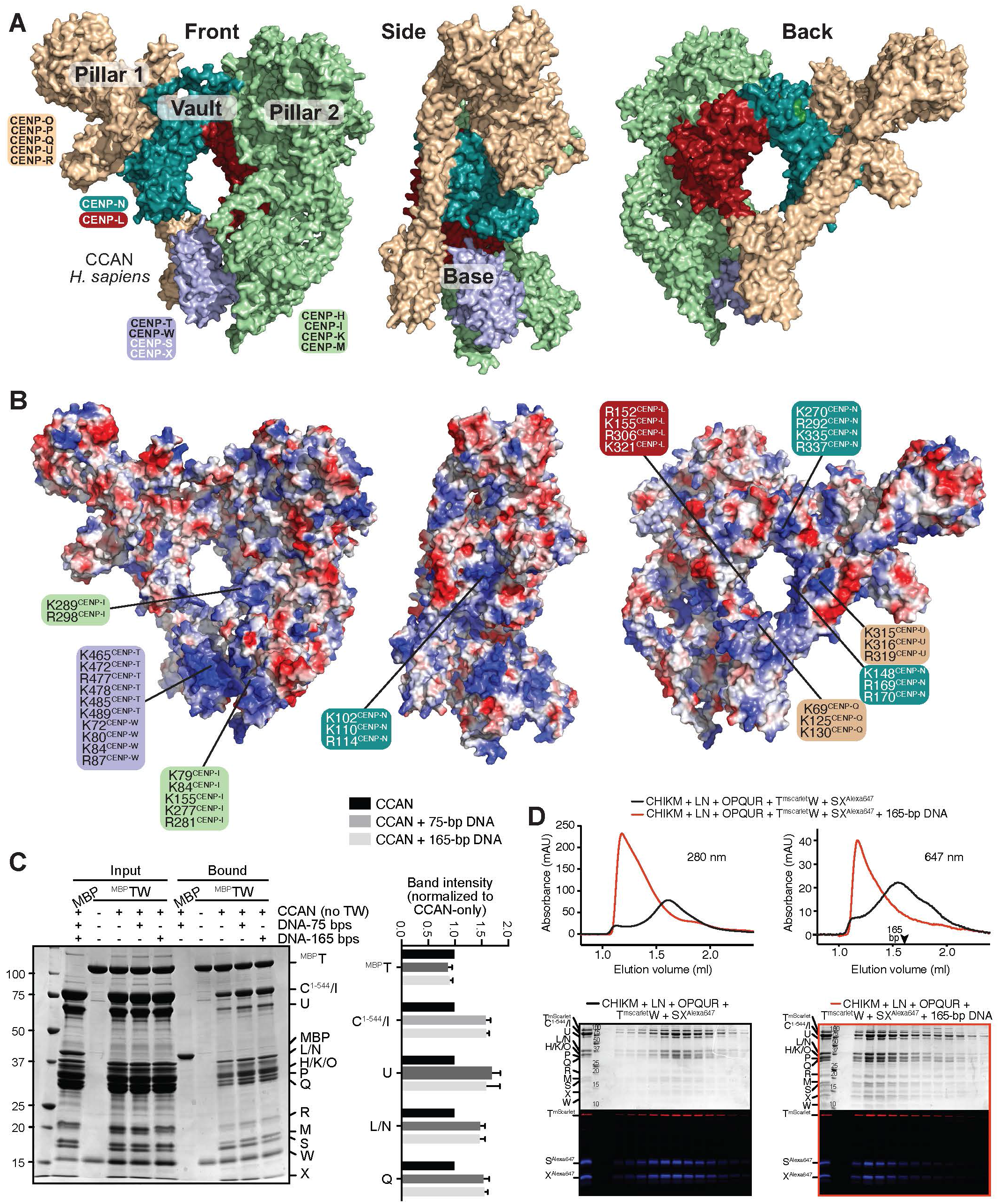
Human CCAN binds DNA. **A**) Three orientations of HsCCAN shown in surface representation. The left panel is from the same view already displayed in Figure 1C. **B**) Surface electrostatics of the HsCCAN complex (red, negative; blue, positive; potential display levels were between - 80 and 80 kT/e) and positively charged residues contributing to potential DNA-binding interfaces discussed in the main text. **C**) *Left*: results from solid phase binding assay on amylose beads, using MBP (negative control) and MBP-CENP-TW as baits. The remaining CCAN subunits were added in solution, with or without the indicated DNA. Beads were recovered by centrifugation, washed, and their content analysed by SDS-PAGE. *Right*: quantified band intensities from three technical replicates normalized to bound CCAN in absence of DNA. Error bars indicate standard deviation. CENP-C/I and CENP-LN comigrated and we report their integrated band intensities. **D**) Size-exclusion chromatography of the indicated complexes with or without 165-bp dsDNA (experiments with 75-bp dsDNA gave analogous results, unpublished observations). Proteins separated by SDS-PAGE were visualized with Coomassie (top) or fluorescence (bottom). Absorbance profiles report on the indicated wavelengths.

To gather evidence for a DNA-binding activity in CENP-16, we immobilized on amylose beads a fusion of CENP-TW with maltose binding protein (MBP), and monitored binding of the remaining CCAN subunits in absence or presence of 75-bp or 165-bp DNA. Both DNAs increased the amounts of the CCAN subunits bound to the immobilized ^MBP^CENP-TW (Figure 3C). Similarly, in size-exclusion chromatography (SEC) experiments, where the elution volume is inversely related to size and elongation, the presence of DNA promoted earlier elution of the CENP-16 CCAN complex and a sharper elution profile, indicative of complex formation and possibly of a stabilization of the complex (Figure 3D). In size-exclusion chromatography experiments with individual CCAN sub-complexes, we found CENP-TW to be a very strong DNA binder (Figure 3 – Supplement 1A), in agreement with previous observations (Nishino et al., 2012; Takeuchi et al., 2014). We have previously reported DNA binding by CENP-HIKM in an EMSA assay at low salt concentrations (Weir et al., 2016), but CENP-HIKM did not appear to bind DNA with sufficient affinity in our SEC assay, nor did CENP-NL (Figure 3 – Supplement 1B-C). Both CENP-C^1-544^HIKM and CENP-OPQUR bound DNA in SEC assays, albeit weakly (Figure 3 – Supplement D-E). Thus, multiple individual CENP-16 sub-complexes interact with DNA, as reported previously for individual complexes (Carroll et al., 2010; Carroll et al., 2009; Nishino et al., 2012; Weir et al., 2016), which predicts a substantial binding affinity of CCAN for DNA. Indeed, the combination of CENP-LN, CENP-C^1-544^HIKM, and CENP-OPQUR in the sample defined as CENP-12 demonstrated strong DNA binding (Figure 3 – Supplement 1F), suggesting additive effects. We anticipate that the different domains of CCAN may form contiguous contacts over a fragment of 65-70 bps of DNA spanning from the front to the back of the complex.

### The CENP-LN vault as a DNA-binding clamp for a single DNA gyre

CENP-L and CENP-N, the proteins that form the vault domain, are paralogs that share a ≈130-residue CENP-LN homology domain (LNHD) and that interact through a distinct C-terminal dimerization domain (Hinshaw and Harrison, 2013; Pentakota et al., 2017) (Figure 4A). The CENP-N^LNHD^ and CENP-L^LNHD^ lean rigidly against pillars 1 and 2, respectively (Figure 1C). Previously reported structures of a CENP-A nucleosome core particle (CENP-A^NCP^) in complex with an N-terminal construct of CENP-N (approximately encompassing residues 1-210 of CENP-N) (Allu et al., 2019; Chittori et al., 2018; Pentakota et al., 2017; Tian et al., 2018) demonstrated that CENP-N^LNHD^ forms extensive interactions with the nucleosome’s DNA, without major contacts with the proteinaceous nucleosome core (Figure 4B, *top two panels*).

**Figure 4.**
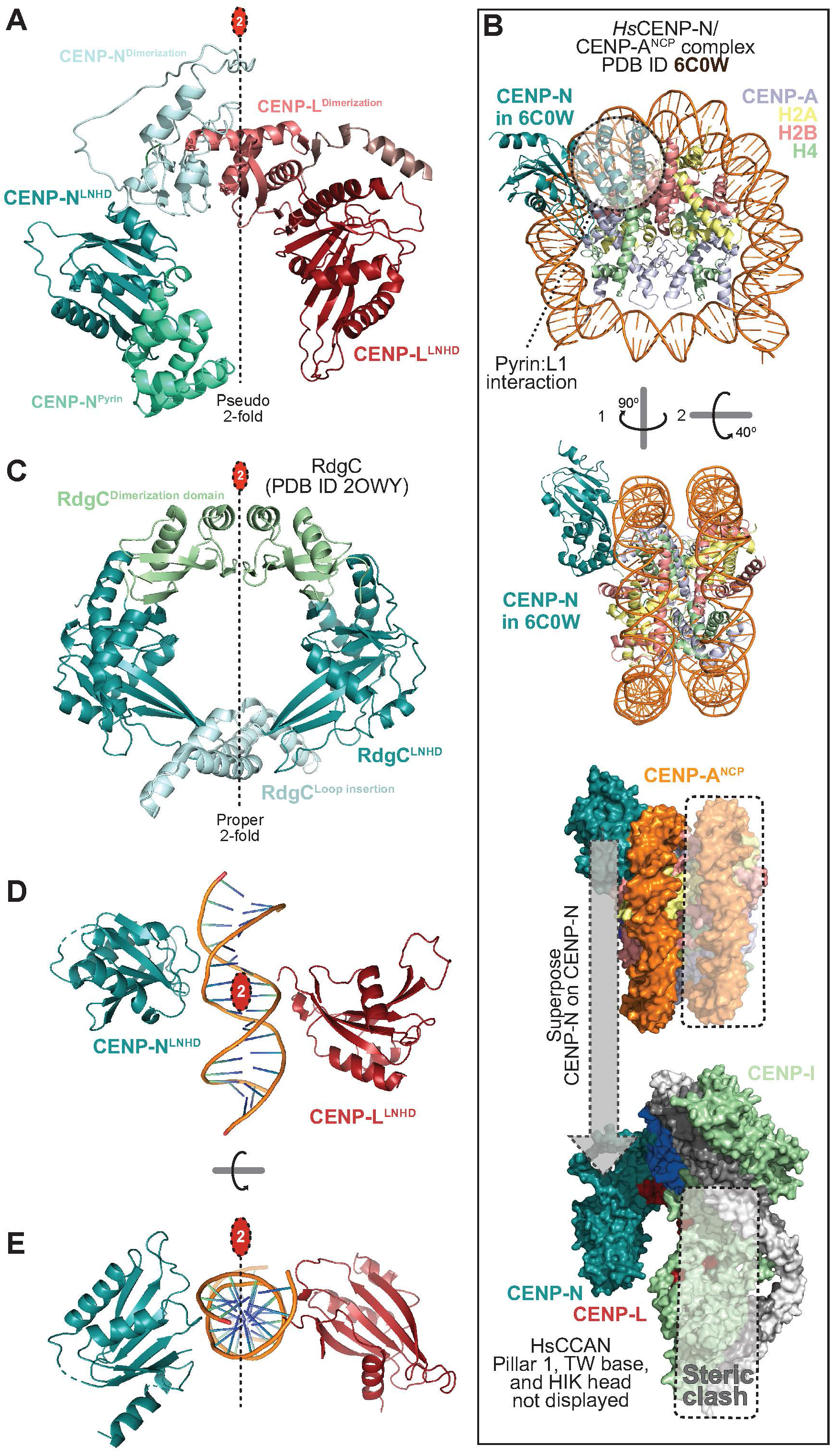
Properties of the CENP-LN vault. **A**) The isolated CENP-LN vault with relevant domains highlighted in different colours. Shown from left to right are the CENP-N pyrin domain (≈80 residues), the CENP-N LNHD (<u>LN</u>-<u>h</u>omology <u>d</u>omain), the dimerization domains of CENP-N and CENP-L, and the CENP-L LNHD. A 2-fold pseudosymmetry axis (interrupted line with red oval at top) relates the CENP-L and CENP-N LNHDs. **B**) *Top*: two rotated cartoon model of the structure of the CENP-N terminal region bound to a CENP-A nucleosome core particle (NCP; PDB ID 6C0W) (Pentakota et al., 2017). CENP-N binds in front of the L1 loop of CENP-A (coloured light blue) and to the DNA. The H2A C-terminal tails were removed to improve clarity. *Bottom*: superposition of CENP-N in 6C0W to CENP-N in the structure of human CCAN predicts a steric clash of the second DNA gyre (boxed) onto subunits of pillar 2 (CENP-HIKM). There is space for a single gyre of DNA in the vault. **C**) Cartoon model of the RdgC homodimer highlighting the LNHDs and other indicated structural elements. **D**) The LNHDs of CENP-LN are highlighted in absence of other structural domain (dimerization, pyrin) and modelled with a 15-bps segment of DNA, extracted from the 6C0W structure shown in panel A and positioned identically relative to CENP-N. The 2-fold pseudosymmetry axis linking the LNHDs coincides with the 2-fold pseudosymmetry axis of the DNA. **E**) A rotated view of the same object displayed in panel F.

Preceding the CENP-N^LNHD^, the 80-residue pyrin domain is also part of the N-terminal region of CENP-N (Figure 4A). In the structures of the CENP-N:CENP-A nucleosome complexes, the CENP-N pyrin domain recognizes the exposed L1 loop of CENP-A (Allu et al., 2019; Chittori et al., 2018; Pentakota et al., 2017; Tian et al., 2018) (Figure 4B). The sequences of CENP-A and H3 diverge significantly in this loop, with important implications for epigenetic centromere inheritance (Black et al., 2004; Black et al., 2007). Mutation of CENP-N residues implicated in L1 loop recognition *in vitro* also prevent CENP-N kinetochore recruitment (Carroll et al., 2009; Chittori et al., 2018; Pentakota et al., 2017), supporting the interaction mode demonstrated by the structural work.

Given this previous evidence, we asked if the nucleosome-binding mode demonstrated with the CENP-N N-terminal region is compatible with CENP-N binding in the context of the entire CCAN complex. To assess this, we superposed the N-terminal regions (pyrin domain and LNHD) of CENP-N in CCAN and in complex with CENP-A^NCP^ (PDB ID 6C0W) and monitored the fit of the nucleosome on the CCAN structure (Figure 4B, *bottom panel*). This demonstrated unequivocally that an octameric CENP-A^NCP^ cannot be accommodated within CENP-16 after superposition of CENP-N, as this would result in a dramatic steric clash of the NCP’s second DNA gyre (i.e. the one distal from CENP-N) with CENP-L and with the HIK head.

This prediction is expected if one considers that our analysis has already identified CENP-LN as being ideally suited to surround a dsDNA filament, but not two adjacent filaments. Indeed, a single dsDNA filament is predicted to fit snugly into the deep, narrow, and positively charged CENP-LN vault (Figure 4 – Supplement 1A). That the vault surrounds a single filament of dsDNA not embedded in a classical nucleosome finds additional support in evolutionary considerations. CENP-L and CENP-N are related to the bacterial protein RdgC, a DNA-binding homodimer that forms a full, closed circle for DNA binding (Ha et al., 2007; Tromer et al., 2019) (Figure 4C). The distant relationship with RdgC seems to imply that CENP-LN originated from the duplication of a DNA binding homo-dimeric singleton (Tromer et al., 2019). This interpretation is further supported by the observation that the CENP-L^LNHD^ and CENP-N^LNHD^ in CENP-16 are related, even in the absence of bound DNA, by a 2-fold pseudosymmetry axis (Figure 4D-E and Figure 4 – Supplement 1A, *right*). With DNA from the proximal gyre of the CENP-N:CENP-A^NCP^ complex modelled in the vault after superposition of CENP-N, the DNA’s own 2-fold pseudosymmetry axis aligns with the 2-fold pseudosymmetry axis of the LNHDs of CENP-L and CENP-N (Figure 4D-E). Thus, CENP-NL in human CCAN is poised to recognize a chromatin structure different from that observed in a classical octameric nucleosome with two adjacent turns of DNA. This conclusion raises a crucial conundrum regarding the effective recognition target of CENP-N. Previous work has implicated CENP-N in specific CENP-A binding at the L1 loop, but our observations now argue that if the CENP-A L1 remains the target of CENP-N in CCAN, it can’t be presented to CENP-N with CENP-A embedded in an octameric nucleosome.

### Nucleosome binding by yeast and human CCAN

An alternative possibility is that CENP-N, at least when incorporated within CCAN, binds only DNA, or that it binds DNA and a different protein target, possibly a different histone-like protein. To shed further light on this question, we continued by analysing the implications for the human complex of a recent structure of the *S. cerevisiae* Ctf19^CCAN^ in complex with a classical octameric Cse4^CENP-A^ nucleosome (PDB ID 6QLD; Figure 5A) (Yan et al., 2019). In this structure, the CENP-LN vault is seen to interact with a loose end of dsDNA unwrapped from the Cse4^NCP^ core (Figure 4 – Supplement 1C-D). Unwrapping of DNA on one side of the nucleosome causes the normally symmetric distribution of DNA on the two sides of the nucleosome to become greatly asymmetric (Figure 5B, *top*, where the red arrows mark the two ends of the DNA, and the vertical bar marks the position of the core’s 2-fold axis). The unwrapped DNA docks in the Chl4^CENP-N^-Iml3^CENP-L^ vault, leaning against Chl4 and making essentially no contacts with Iml3 or more generally with pillar 2 (Figure 5C-D and Figure 4 – Supplement 1C-D). The position of Chl4^CENP-N^ and of CENP-N relative to the NCP in the Ctf19^CCAN^:NCP and CENP-N:CENP-A^NCP^ structures (PDB ID 6C0W) are therefore entirely unrelated, to the point that in the yeast structure CENP-N does not contact the L1 loop of CENP-A at all, contrary to the human CENP-N:CENP-A^NCP^ structure, where CENP-N directly faces the L1 loop (Figure 5B). In fact, there are almost no visible contacts of Ctf19^CCAN^ and the proteinaceous Cse4^CENP-A^ nucleosome core in PDB ID 6QLD. This perplexing observation suggests that Ctf19^CCAN^ connects to the centromeric nucleosome almost exclusively through the intrinsically disordered segments of the (largely invisible) Mif2^CENP-C^ “blueprint”. How Mif2 interacts with yeast CCAN remain elusive.

**Figure 5.**
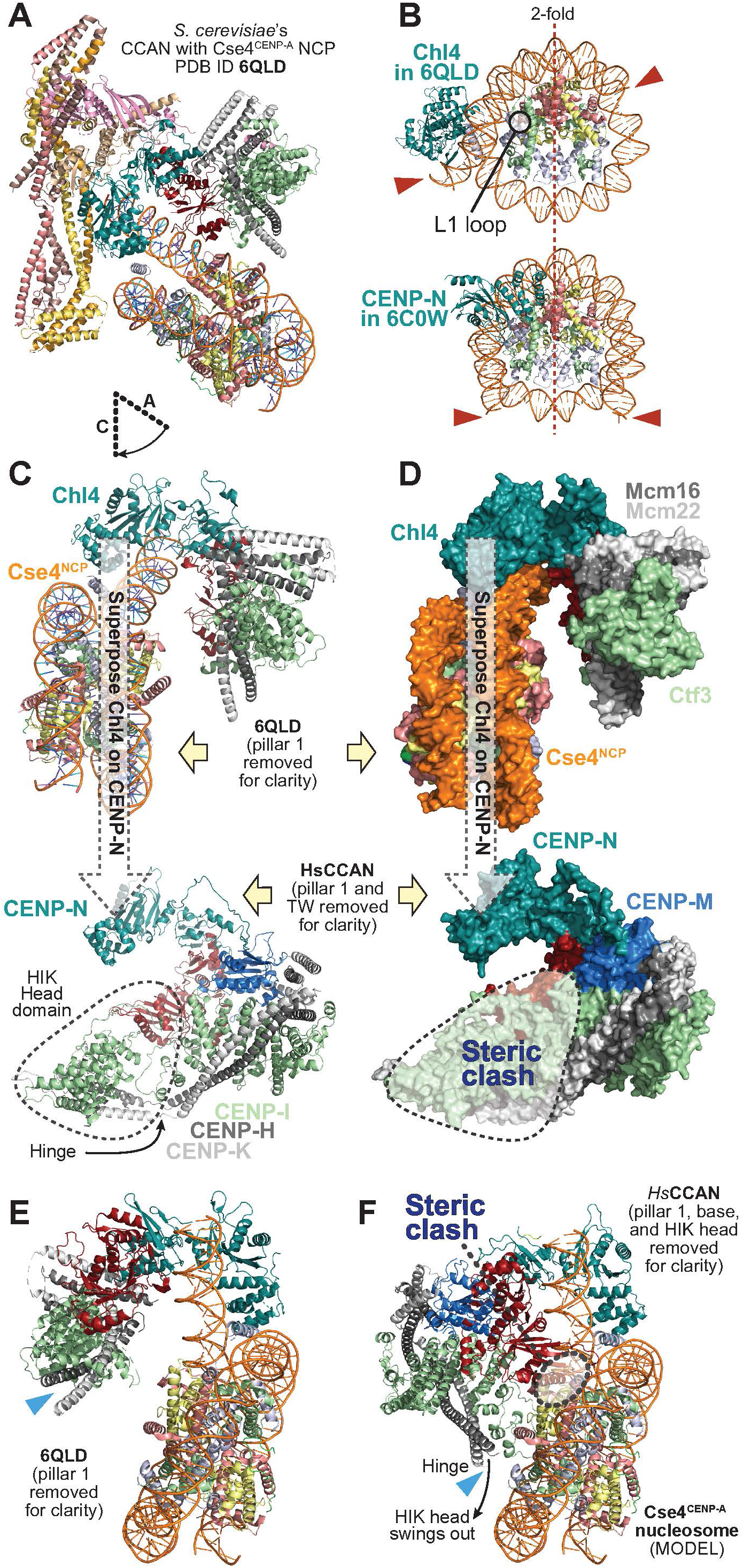
A yeast Ctf19^CCAN^:CENP-A nucleosome structure is a poor model for the human CCAN. **A**) Cartoon view of the *S. cerevisiae* Ctf19^CCAN^:CENP-A nucleosome complex (PDB ID 6QLD) (Yan et al., 2019). **B**) Comparison of CENP-N on the yeast and human nucleosome structures demonstrates a different position. The protein cores of the nucleosomes have the same orientation, as shown by the indicated position of the 2-fold pseudosymmetry axis. Red arrows point to the DNA ends in the two structures, which are different. They are offset in PDB ID 6QLD, due to the unwrapping of the nucleosomal DNA attracted inside the vault. They are instead roughly symmetric around the 2-fold pseudosymmetry axis in PDB ID 6C0W. The H2A C-terminal tails were removed to improve clarity. **C**) *Top*: A rotated view of the yeast complex (PDB ID 6QLD) already shown in panel A. The rotation between panels A and C is schematized. Pillar 1 was removed to improve clarity. The position of the Chl4^CENP-N^ was used to superpose the structure to the human CCAN complex (*bottom*, pillar 1 and the base were also removed for clarity, so that only the vault and pillar 2 are shown). The HIK Head domain is boxed. **D**) The same objects shown in panel C are now shown as surfaces. Superposition of CENP-N^Chl4^ predicts a dramatic steric clash of the nucleosome with the HIK head domain, indicating that the yeast nucleosome structure is not a good predictor for the complex of human CCAN with a CENP-A nucleosome. **E**) PDB ID 6QLD is displayed as in panel C, but rotated approximately 180°. There are no contacts of pillar 2 with the Cse4^NCP^. **F**) Human CCAN was modelled onto PDB ID 6QLD by aligning CENP-N on Chl4 as in panel D. The predicted steric clash of HIK head with the Cse4^CENP-A^ nucleosome could be solved if the HIK head swung out by rotation about the hinge. There are residual predicted clashes of CENP-L with H2A:H2B. Due to the different direction of pillar 2 relative to CENP-N^Chl4^ in the yeast and human complexes, pillar 2 adopts a very different orientation

We asked if the structure of the Ctf19^CCAN^:Cse4^CENP-A^ nucleosome complex provides a plausible model for the human complex. To assess this, we superposed Chl4^CENP-N^ in 6QLD with CENP-N in our structure of human CENP-16, and observed if the nucleosome in PDB ID 6QLD could be accommodated in the human structure without major steric clashes. This procedure evidenced a dramatic clash of the yeast nucleosome with the HIK head of human CCAN (Figure 5D, *bottom*), with an additional but less dramatic predicted overlap between CENP-L and one of the two H2A:H2B dimers. The main clash could be resolved through a hypothetical rotation of the HIK head about a hinge (Figure 5C, *bottom*) and causing the HIK head to be swung out of its observed position, similar to the yeast Ctf19^CCAN^, where the HIK head undergoes an approximately 90° rotation upon binding to the nucleosome (Yan et al., 2019). Due to fundamental differences in the respective orientation of pillar 2 relative to CENP-N (Figure 1E), however, the yeast complex (PDB ID 6QLD) and the human model built on it would appear quite different even after removing the clash with the HIK head (Figure 5E-F): in the Ctf19^CCAN^:Cse4^CENP-A^ complex (PDB ID 6QLD) pillar 2 does not even contact the Cse4^CENP-A^ nucleosome (Figure 5C,E). Conversely, in the modelled human complex, pillar 2 is predicted to make multiple contacts with the nucleosome (Figure 5F). Thus, CCAN may bind an octameric CENP-A nucleosome positioned as in the structure of the yeast Ctf19^CCAN^-Cse4^NCP^ complex (PDB ID 6QLD), but to avoid predicted steric clashes a very large rotation of the HIK head about a hinge point (indicated by blue arrowheads in Figure 5E-F), as well as other less significant changes around CENP-L, would have to be enabled. The way pillar 2 and the nucleosome interact in PDB ID 6QLD and in the resulting human structure would still differ substantially, a consequence of the different opening of the vault.

### Open and closed vaults in point and regional centromeres

The different organization of human and yeast CCAN^Ctf19^ might have multiple causes, including the stabilizing presence of CENP-M in the human complex but not in the yeast complex, and a much larger interface of pillar 1 with the vault in the human complex relative to the yeast complex (Figure 2 – Supplement 1B). A superposition of CENP-N with Chl4^CENP-N^ shows a reciprocal rotation within the Iml3^CENP-L^:Chl4^CENP-N^ dimer that causes the vault to be much shallower and open in comparison to CENP-LN (Figure 6A). As a consequence, the 2-fold pseudosymmetry of the LNHDs identified in the human complex with a modelled DNA (Figure 4D-E and Figure 4 – Supplement 1A) is broken in the yeast Ctf19^CCAN^:Cse4^CENP-A^ nucleosome complex (PDB ID 6QLD; Figure 4 – Supplement 1B-D). Modelling with AF2 of Iml3:Chl4 orthologs in yeast species featuring point centromeres and in diverse organisms with regional centromeres demonstrated that the complexes cluster into two different groups, with complexes from organisms with point centromeres showing shallow vaults and complexes from species with regional centromeres showing deep vaults (Figure 6B). This result implies that at least in part, the structural properties of the vault are moulded by intrinsic sequence features, in addition to influences from its embedding in the CCAN^Ctf19^ complex. Thus, the different structural features of CCAN^Ctf19^ complexes in point and regional centromeres are conserved, perhaps hinting to a different composition of the fully assembled kinetochore complexes.

**Figure 6.**
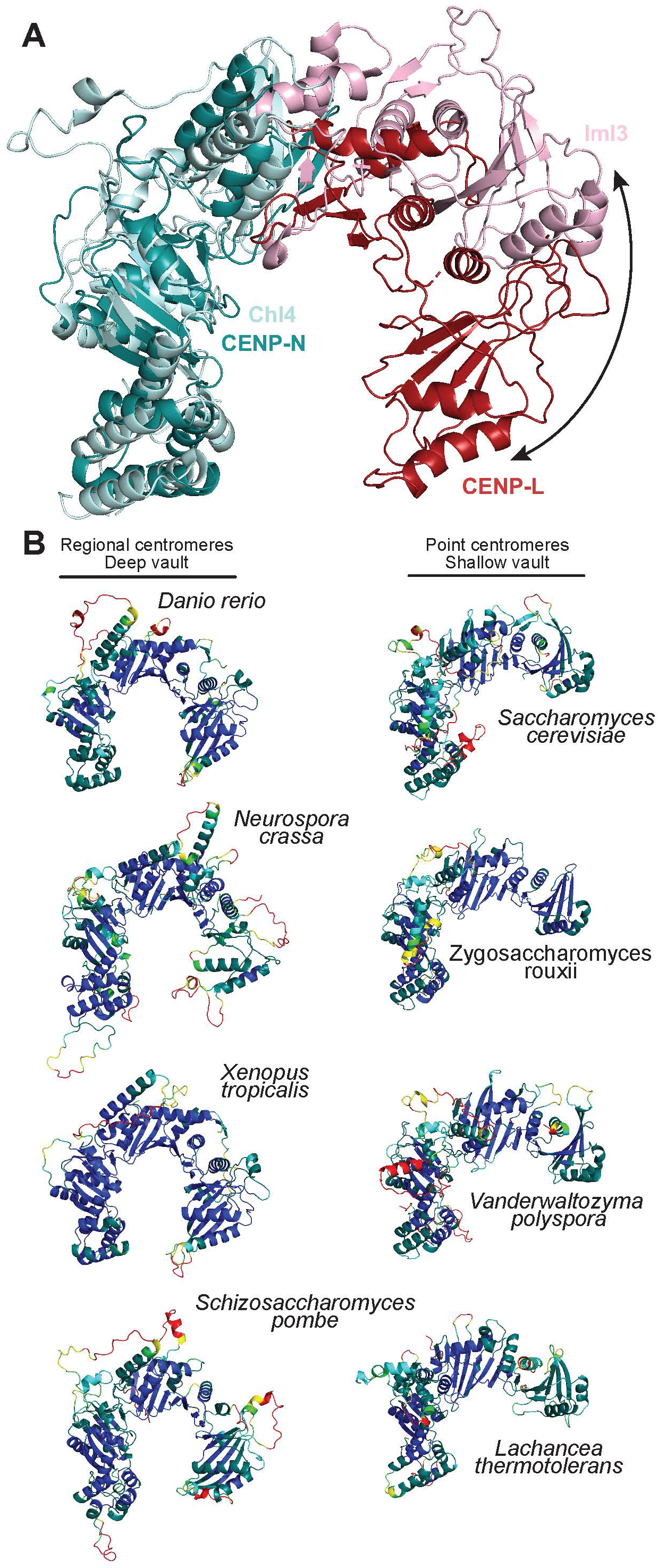
Deep and shallow vaults in CENP-LN orthologs. **A**) Superposition of CENP-N and Chl4 in human CENP-LN (respectively in deepteal and firebrick) and Iml3^CENP-L^:Chl4^CENP-N^ (respectively in cyan and pink). The curved arrow emphasizes the relative rotation of CENP-L about a hinge point in the dimerization domain. **B**) Gallery of AF2 predictions of vertebrate and yeast CENP-LN orthologs from the indicated species, represented in pLDDT score (blue, high confidence prediction; red, low confidence prediction).

## Discussion

The structure of human CCAN is a milestone in the study of the organization of centromeric chromatin. It builds on early proteomic analyses that identified the majority of vertebrate CCAN subunits (Foltz et al., 2006; Izuta et al., 2006; Obuse et al., 2004; Okada et al., 2006) and on subsequent work of biochemical reconstitution and structural analysis (Ali-Ahmad et al., 2019; Allu et al., 2019; Ariyoshi et al., 2021; Carroll et al., 2010; Carroll et al., 2009; Guo et al., 2017; Kato et al., 2013; McKinley et al., 2015; Pentakota et al., 2017; Pesenti et al., 2018; Walstein et al., 2021; Weir et al., 2016). The structure and biochemical work reported here demonstrates the overall organization of human CCAN, with insights on unique subunits, including CENP-M and CENP-R, and fundamental differences with the yeast complex. Our work also provides the best demonstration so far that regional centromeres are assembled from the repetition of an individual structural module related to that of the *Saccharomyces cerevisiae* Ctf19^CCAN^ complex (Hinshaw and Harrison, 2019, 2020; Yan et al., 2019; Zhang et al., 2020). Regretfully, we have been unable to obtain a sufficiently homogeneous complex of CCAN with DNA or nucleosomes for structural analysis (unpublished results), a priority for our future studies. Nevertheless, the structure of human CCAN has important implications for the basic concepts that have accompanied the vertebrate centromere field in recent years.

The most unexpected observation concerns CENP-N, initially identified with CENP-C as the only direct and specific binder of CENP-A nucleosomes (Carroll et al., 2010; Carroll et al., 2009). The structure of human CCAN shows that the CENP-LN dimer is ideally suited to surround a single filament of dsDNA, as also supported by a suggestive 2-fold pseudosymmetry arrangement of the LNHDs of CENP-L and CENP-N in the vault. As the CENP-LN complex alone does not have high affinity for DNA, its interaction with DNA may be eminently topological. If the CENP-LN vault is designed to bind a single filament of dsDNA, a most pressing question is whether the pyrin domain of CENP-N, when embedded in CCAN, can decode the CENP-A L1 loop as implied by previous structural work with isolated CENP-N deletion constructs bound to CENP-A nucleosomes (Allu et al., 2019; Chittori et al., 2018; Pentakota et al., 2017; Tian et al., 2018). Based on 1) direct evidence from the structure of the yeast Ctf19^CCAN^:Cse4^CENP-A^ nucleosome complex (PDB ID 6QLD), where the Chl4^CENP-N^ pyrin domain is mainly engaged in DNA binding and does not even face the L1 loop; and 2) indirect evidence from our own work showing that it is not possible to fit an octameric CENP-A nucleosome into the human CCAN structure while concomitantly satisfying the condition that the CENP-N pyrin domain faces the CENP-A L1 loop (Allu et al., 2019; Chittori et al., 2018; Pentakota et al., 2017; Tian et al., 2018), the most immediate answer is ‘no’.

If CENP-N does not contribute to the specificity of CENP-A recognition, only CENP-C, to the best of our knowledge the only additional specific CENP-A binder in CCAN, remains to explain it. The implications of this conclusion are summarized in Figure 7A, which incorporates related features of the human and yeast CCAN^Ctf19^. The dsDNA filament that occupies the vault of CCAN is connected to an octameric CENP-A nucleosome (or a dinucleosome (Walstein et al., 2021)). It enters CCAN from the front (like in the yeast Ctf19^CCAN^:Cse4^CENP-A^ nucleosome structure, PDB ID 6QLD) (Yan et al., 2019) or from the back. In the “swung-in” conformation of the human complex, CENP-TW could be expected to bind and stabilize the dsDNA filament as the “base” under the vault. A possible position of the CENP-A nucleosome is inferred from the structure of the Ctf19^CCAN^:Cse4^CENP-A^ nucleosome complex (PDB ID 6QLD) (Yan et al., 2019). If the nucleosome positioned itself in the human complex as in yeast, an intolerable steric clash with the “swung-in” HIK head and base would arise where the complete histone core would have to be replaced by the HIK head (Figure 5C-D). This could be resolved by 1) repositioning the nucleosome in a very different position, or 2) adopting a swung-out conformation of the HIK head and CENP-TW base, making space for a nucleosome positioned like in yeast and allowing CENP-TW to bind linker DNA, which it prefers over nucleosome-wrapped DNA (Takeuchi et al., 2014). With a swung-out HIK head, however, the details of the interaction with the CENP-A^Cse4^ nucleosome in yeast and humans would still differ substantially due to the different angle of divergence of pillar 2 (Figure 5E-F).

**Figure 7.**
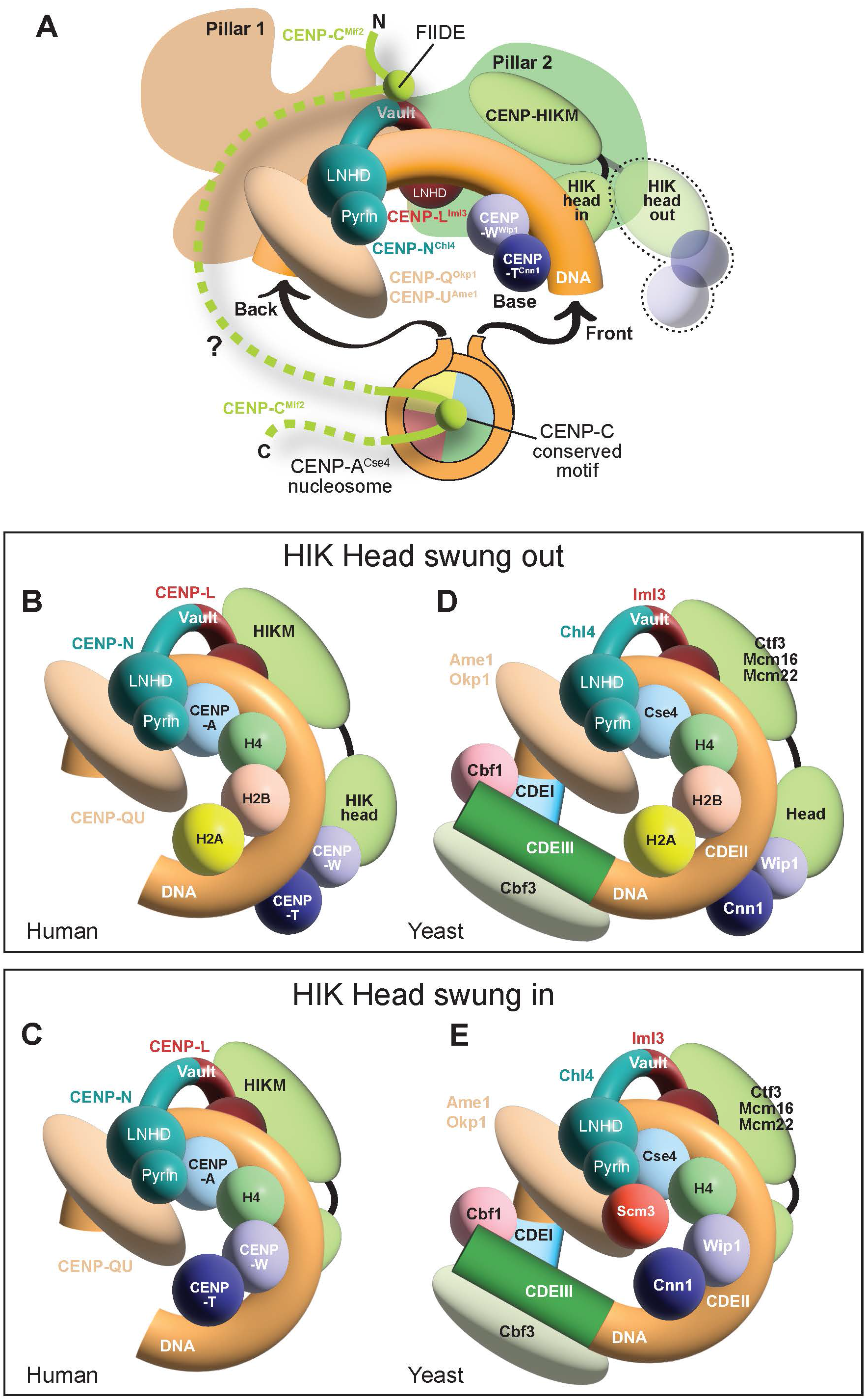
Models of centromere:chromatin interaction. **A**) Common features of the CCAN^Ctf19^:CENP-A^Cse4^ interaction in yeast and humans. The CENP-L^Iml3^N^Chl4^ vault is occupied by dsDNA that emerges from a CENP-A^Cse4^ nucleosome that is otherwise not directly integrated in CCAN ^Ctf19^ and only connected to it through CENP-C^Mif2^, which acts as the crucial link between the nucleosome and the CCAN^Ctf19^. CENP-C^MIf2^ is flexible (dotted line), enabling multiple binding modes observed or predicted in yeast and humans. The conserved motif binds the CENP-A^Cse4^ nucleosome. The FIIDE motif is only detected in human CENP-C, but an equivalent region of Mif2^CENP-C^ binds to Iml3^CENP-L^:Chl4^CENP-N^ in *S. cerevisiae* (Hinshaw and Harrison, 2013). In the “swung-in” conformation the HID head positions CENP-T^Cnn1^W^Wip1^ in the observed “base” position, which can only be reached in the human complex due to the higher divergence of pillar 2 in the yeast complex. A “swung-out” conformation is also shown. **B**) In this hemisome model CENP-A:H4 faces the pyrin domain of CENP-N. A single filament of dsDNA is allowed inside the CENP-LN vault. The CENP-TW base connected to the HIK-head is in a swung-out conformation that permits an interaction of CENP-A:H4 with H2A:H2B, with which CENP-TW would otherwise crash. CENP-QU may contribute, alone or in complex with other proteins, to prevent CENP-A dimerization. **C**) The same hemisome complex, but with H2A:H2B replaced by CENP-TW as expected for the “swung-in” conformation observed in our structures. **D**) The CDEII core of the yeast point centromere is ≈85-bps-long, and flanked by CDEI and CDEIII motif that associate with Cbf1 and Cbf3. A hemisome model has been proposed for this organism (see main text). **E**) Another hemisome model may explain depletion of H2A:H2B at yeast centromeres as well as a function of Scm3 in preventing Cse4^CENP-A^ dimerization.

Emphasis on predicted differences between the human and yeast complexes aims to expose a crucial aspect of the model(s) in Figure 7A. The structure of the complex of Ctf19^CCAN^ with the Cse4^CENP-A^ nucleosome (PDB ID 6QLD) (Yan et al., 2019) identified too few contacts to account for the exquisite specificity expected of this interaction. CENP-C^Mif2^, in the absence of CENP-N the only expected generator of specificity in the interaction of CCAN with CENP-A, is predicted to be mainly intrinsically disordered. It may therefore connect CCAN^Ctf19^ and the CENP-A^Cse4^ nucleosome with substantial flexibility and conformational freedom. The resulting complexes may be “fuzzy” rather than rigidly sculpted. In such a framework, the predicted and quite substantial differences between the human and yeast complexes discussed above may all be encompassed in the model in Figure 7A and be ultimately compatible with function.

Mif2f^CENP-C^ is for the most part invisible in the yeast Ctf19^CCAN^:Cse4^CENP-A^ nucleosome complex (PDB ID 6QLD) (Yan et al., 2019) (Figure 7A). The only visible segment of Mif2f^CENP-C^ is the conserved motif (residues 284-305), shown to bind the acidic patch of H2A:H2B and the CENP-A C-terminal tail (Yan et al., 2019), a binding mode previously demonstrated in vertebrates (Ariyoshi et al., 2021; Kato et al., 2013; Yan et al., 2019). Mif2^CENP-C^ is also known to interact with Iml3^CENP-L^:Chl4^CENP-N^(Hinshaw and Harrison, 2013), an interaction that is conserved in humans, where additional interactions of CENP-C with CENP-HIKM have also been observed (Hinshaw and Harrison, 2013; Hornung et al., 2014; Klare et al., 2015; McKinley et al., 2015; Nagpal et al., 2015; Pentakota et al., 2017). The specific motifs involved in these interactions with CCAN^Ctf19^ may not be fully conserved: the 303-FIIDE-307 CENP-C motif we describe, previously shown to be instrumental for the recruitment of CENP-LN to the kinetochore (Pentakota et al., 2017), is not recognizable in Mif2^CENP-C^.

The dismissal of CENP-N as a specificity factor for CENP-A recognition should not be taken light-heartedly. Mutations of the CENP-N region in the pyrin domain involved in L1 binding ablate CENP-A binding *in vitro* and CENP-N localization *in vivo* (Allu et al., 2019; Carroll et al., 2010; Carroll et al., 2009; Chittori et al., 2018; Pentakota et al., 2017; Tian et al., 2018). If the structure of the yeast Ctf19^CCAN^:Cse4^CENP-A^ nucleosome complex did not confirm Chl4^CENP-N^ as L1 loop decoder, the structure of human CCAN now at hand prompts us to identify what requirements should be met for the CENP-N pyrin domain to recognize the L1 loop. Two crucial conditions emerge. First, CENP-A:H4 ought to be presented to CENP-LN in a structure different from a classical 2-turn nucleosome, as only the CENP-N-proximal filament of dsDNA, with its associated CENP-A:H4, would be allowed in the CENP-LN vault without major steric clashes. Second, homo-dimerization of CENP-A would likely be prevented (Black et al., 2004; Tachiwana et al., 2011), due to a predicted steric clash with the CENP-QU N-terminal domain, contiguous to the CCAN base at the back of the CENP-LN tunnel.

A structure that satisfies these two conditions is the so-called hemisome, a half-nucleosome sequence of histones with the order CENP-A:H4:H2B:H2A (Figure 7B). In CCAN, H2A:H2B could be part of the hemisome only if the HIK head and CENP-TW base adopted a swung-out conformation. In the swung-in conformation of our structure, H2A:H2B would be excluded due to a dramatic steric clash with CENP-TW, which could then pair with CENP-A:H4 (Figure 7C). Models invoking a hemisome as in Figure 7B-C are entirely speculative at this time, and may also seem to impose prohibitive assumptions. There is unquestionable evidence that CENP-A assembles *in vitro* into classic octameric nucleosomes with adjacent DNA gyres, and that classical octameric nucleosomes are also the predominant form of CENP-A isolated from chromatin after extensive nuclease treatment (Black and Cleveland, 2011; Dunleavy et al., 2013; Hasson et al., 2013; Nechemia-Arbely et al., 2017; Tachiwana et al., 2011). As discussed in the Introduction, however, extensive nuclease treatments may promote the disassembly of centromeric chromatin (Ando et al., 2002), raising the question whether previous studies might have neglected a handful of CENP-A molecules embedded in a different and possibly larger (than a mononucleosome) chromatin structure (see also (Walstein et al., 2021)).

How can the alternative models discussed in Figure 7A-C be tested? Reconstitution experiments exposing CCAN^Ctf19^ to a pre-assembled and greatly stable octameric CENP-A nucleosome will with some likelihood promote assembly of a complex with sufficient stability for structural analysis. This will not necessarily prove that the resulting structure, aligned with the model in Figure 7A, is physiologically relevant. The same approach would not be helpful towards testing the speculative models discussed in Figure 7B-C, as they predict a different type of nucleosome organization that may need to be stabilized differently from classical nucleosomes. Two crucial stabilizing factors of octameric nucleosomes are the dimerization of CENP-A (or H3) and the stabilization of the left-handed helical staircase of histone octamers by the H2A C-terminal docking domain (Eickbush et al., 1988; Shukla et al., 2011). Neither would be at stake in the hemisome structure postulated in Figure 7B. In the model in Figure 7C, also a hemisome model but with CENP-TW replacing H2A:H2B, the C-terminal extension of CENP-T is buried at the interface with HIK (this study and (Hinshaw and Harrison, 2020; Zhang et al., 2020)). These alternative structures will have much reduced stability of their own compared to octameric nucleosomes, and *ad hoc* procedures for their incorporation in, and stabilization by, CCAN may be necessary for successful reconstitution. Enzymes overcoming kinetic barriers large enough to slow down spontaneous execution may even be required. Be that as it may, our attempts at reconstituting a CCAN:chromatin structure with DNA, CENP-TW (with or without H2A:H2B) and CENP-A:H4 bound to the CENP-NL vault were hitherto unsuccessful (unpublished results). Furthermore, we find that at least the hemisome model in Figure 7C is not supported by predictions of AF2 and related programs (unpublished results). Thus, we cannot claim own experimental evidence supporting models like those in Figure 7B-C. Yet, we cannot rule them out, as we might have failed to identify appropriate conditions for their assembly.

These concerns are especially relevant to interpret recent work on the Ctf19^CCAN^ complex and Cse4^CENP-A^ nucleosome of *S. cerevisiae*. In this organism, Cse4^CENP-A^ resides on a central DNA core of 78-86 bps named CDEII (Camahort et al., 2009; Furuyama and Biggins, 2007; Henikoff et al., 2014; Keith and Fitzgerald-Hayes, 2000; Krassovsky et al., 2012; Meluh et al., 1998). Flanking CDEII, two additional regions of 8 and ≈25 bps, known respectively as CDEI and CDEIII, bind the additional factors Cbf1 and Cbf3 complex. The question how the Ctf19^CCAN^ complex relates to the centromeric DNA of *S. cerevisiae* should be considered in the context of these fundamental specificities, something that the work so far, limited to a complex with an octameric nucleosome on a 147 bps Widom 601 DNA sequence (PDB ID 6QLD) (Yan et al., 2019), has not yet fully achieved. Cogently, given the small size of CDEII, a centromeric nucleosome with two full turns seems unlikely. A Cse4^CENP-A^:H4:H2B:H2A hemisome has been proposed as a possible alternative (Dalal et al., 2007; Furuyama et al., 2013; Henikoff et al., 2014; Talbert and Henikoff, 2020). This hemisome model, illustrated in Figure 7D, has been extensively debated in the last fifteen years (Black and Cleveland, 2011; Dalal et al., 2007; Dunleavy et al., 2013). It remains highly conjectural, but was shown to neatly explain the pattern of H4 S47C-anchored cleavage mapping at *S. cerevisiae* centromeres (Henikoff et al., 2014). We note that the hemisome model illustrated in Figure 7D is closely reminiscent of the speculative model of human CCAN discussed in Figure 7B, a model that we developed with the question in mind whether specific recognition of CENP-A by CENP-N is compatible with CCAN structure. The coincidence is remarkable and worth reporting, even though its significance is currently unclear.

Further complicating the picture, histone H2A and H2B have been shown to be depleted from centromeres both in *S. cerevisiae* and *S. pombe* (Mizuguchi et al., 2007; Rossi et al., 2021; Williams et al., 2009; Xiao et al., 2011), opposing observations notwithstanding (Krassovsky et al., 2012; Pinto and Winston, 2000; Westermann et al., 2003). At yeast centromeres, the possible depletion of H2A:H2B had been discussed in the context of evidence supporting the existence of a hexasome of Cse4^CENP-A^:H4 with Scm3. Scm3 is a Cse4 chaperone and a stable centromere resident at all cell cycle stages in *S. cerevisiae* (Mizuguchi et al., 2007; Xiao et al., 2011). Subsequent structural work indicated that Scm3 binds the Cse4^CENP-A^ dimerization interface and competes with dimerization (Cho and Harrison, 2011; Dechassa et al., 2011), questioning the hexasome model, which assumed the dimerization of Cse4^CENP-A^:H4 in a tetrasome. A speculative alternative explanation is that Scm3, in addition to depositing Cse4^CENP-A^, stably suppresses Cse4^CENP-A^ dimerization, an expected and potentially beneficial function if the basic structure of yeast centromeres were a hemisome. How could the depletion of H2A:H2B be accounted for, however? Our speculative model in Figure 7C, where H2A:H2B are replaced with CENP-TW, may serve as inspiration to answer this question. H2A:H2B may be replaced with Cnn1^CENP-T^:Wip1^CENP-W^ if a “swung-in” conformation of the yeast complex, similar to that observed in humans, were possible (Figure 7E). We predict, however, that this would require a more complex restructuring of pillar 2 towards the conformation it adopts in human CCAN.

In conclusion, our model of human CCAN is in principle compatible with two classes of models for centromeric chromatin. In one class (exemplified by Figure 7A), which is more strongly supported by current experimental evidence but far from being established, the CCAN assembles on dsDNA and flanks a regular octameric CENP-A^Cse4^ nucleosome, to which it connects specifically mainly or exclusively through CENP-C^Mif2^ (Hinshaw and Harrison, 2013). In the other class (exemplified by Figure 7B-E), CENP-N^Chl4^, in addition to CENP-C^Mif2^, contributes to the specificity of the CCAN^Ctf19^:CENP-A^Cse4^ nucleosome interaction by interacting with the L1 loop of CENP-A^Cse4^. In this second class of models, the CENP-A chromatin interacting with CCAN cannot be in the form of a regular octameric nucleosome, while hemisomes may be expected. If this alternative chromatin exists at all, its reconstitution may be considerably more challenging in view of different, and currently unknown, stabilization requirements. Future work will have to address systematically the implications and value of these models *in vitro* and *in vivo*, testing them with a combination of biochemical reconstitution, structural analysis, and mutational validation.

## Materials and Methods

### Plasmids

Plasmids to express recombinant CENP-LN, -OPQUR, -TWSX, -TW, -SX complexes and CENP-A containing nucleosomes were generated as previously described (Pentakota et al., 2017; Pesenti et al., 2018; Walstein et al., 2021). To generate plasmid for expressing CENP-C^1-544^HIKM in insect cells, plasmid to express N-terminus 6xHis tagged CENP-C^1-544^ was generated by Gibson cloning method, codon optimized cDNA of CENP-C^1-544^ (GeneArt, Life Technologies) was inserted in a modified pLIB vectors containing sequences for the 6xHis followed by TEV protease. The codon optimized cDNA of CENP-I, -H, -K- and -M were inserted by Gibson cloning method in to unmodified pLIB plasmids. These pLIB plasmids generated so-forth were used to insert the CENP-H, -K, -I, -M and 6His-CENP-C^1-544^ sequences into a baculovirus-based multigene-expressing vectors, pBIGa (Weissmann et al., 2016) by Gibson assembly. To generate plasmids for expressing ^MBP^CENP-TW and ^mScarlet^CENP-TW in bacterial cells, codon optimized cDNA of CENP-T, and -W were inserted by Gibson in a pETDuet plasmid containing a ^6His-TEV^CENP-W and a ^Halo-TEV^CENP-T (Walstein et al., 2021) to generate plasmids co-expressing the following constructs: ^6His-TEV^CENP-W/^MBP^CENP-T and ^6His-TEV^CENP-W/^mScarlet^CENP-T. The plasmids pUC18 containing 165-bps and 75-bps CEN1 (centromere 1) like sequences GTGGTAGAATAGGAAATATCTTCCTATAGAAACTAGACAGAATGATTCTCAGAAACTCCTTTGTGATGTGTGCGTTCAACTCACAGAGTTTAACCTTTCTTTTCATAGAGCAGTTAGGAAACACTCTGTTTGTAATGTCTGCAAGTGGATATTCAGACGCCCTTG and ATCCGTGGTAGAATAGGAAATATCTTCCTATAGAAACTAGACAGAATGATTCTCAGAAACTCCTTTGTGATGGAT were generated as previously described (Walstein et al., 2021).

### Protein expression and purification

CENP-LN, -OPQUR, -TWSX and -SX complexes were expressed and purified according to previously published protocols (Pentakota et al., 2017; Pesenti et al., 2018; Walstein et al., 2021). Expression and purification of CENP-C^1-544^HIKM was performed following a protocol adapted from (Klare et al., 2015). TnAo38 cells infected with a virus:culture ratio of 1:50 and incubated for 72 h at 27 °C. Cell pellets were harvested, washed in 1× PBS, and finally resuspended in a buffer containing 50 mM HEPES 7.0, 500 mM NaCl, 5 mM MgCl_2_, 5% glycerol, 10 mM imidazole, 2 mM TCEP, 0.2 mM PMSF, and 10 µg/ml DNase. Cells were lysed by sonication, and cleared for 1 h at 100,000g. Cleared cell lysate was then applied over a 5 ml HisTrap FF column (Cytiva) and washed first with Washing buffer (50 mM HEPES 7.0, 500 mM NaCl, 5 mM MgCl_2_, 5% glycerol, 10 mM imidazole, and 2 mM TCEP), secondly with High salt washing buffer (50 mM HEPES 7.0, 1 M NaCl, 5 mM MgCl_2_, 5% glycerol, 10 mM imidazole, and 2 mM TCEP) followed by Washing buffer again and thirdly with Imidazole washing buffer (50 mM HEPES 7.0, 500 mM NaCl, 5 mM MgCl_2_, 5% glycerol, 40 mM imidazole, and 2 mM TCEP). CENP-C^1-544^HIKM complex was eluted with Elution buffer (50 mM HEPES 7.0, 500 mM NaCl, 5 mM MgCl_2_, 5% glycerol, 200 mM imidazole, and 2 mM TCEP). The fractions containing CENP-C^1-544^HIKM were pooled, and the His tag cleaved overnight at 4 °C with TEV protease (in house production). CENP-C^1-544^HIKM in solution was then adjusted to a salt concentration of 300 mM, before loading on a 5 ml HiTrap Heparin HP column (Cytiva), equilibrated in 20 mM HEPES pH 7.0, 300 mM NaCl, 5% glycerol, 2 mM TCEP. Bound proteins were eluted with a gradient of 300-1000 mM NaCl over 30 column volumes, and peak fractions corresponding to CENP-C^1-544^HIKM were pooled and concentrated in a 50 kDa MW Amicon concentrator (Millipore). CENP-C^1-544^HIKM was then loaded onto a Superose 6 16/600 (Cytiva) in 20 mM HEPES 7.0, 500 mM NaCl, 5% glycerol, 3 mM TCEP. The sample was concentrated and flash frozen in liquid N2 before use. ^MBP^CENP-TW and ^mScarlet^TW complexes were purified using the previously described protocol for CENP-TW wild type (Walstein et al., 2021). Preparations of CENP-A-containing nucleosomes were carried out as described (Walstein et al., 2021) modified from the previously published protocol (Guse et al., 2012).

### Recombinant proteins fluorescence labeling

CENP-SX was labeled using Alexa Fluor 647 Protein Labeling Kit (ThermoFisher Scientific, Waltham, US-MA) according to the manufacturer instructions.

### Analytical SEC

Analytical size exclusion chromatography was carried out on a Superose 6 5/150 (Cytiva, Marlborough, US-MA) in a buffer containing 20 mM HEPES pH 6.8, 300 mM NaCl, 2.5 % (v/v) glycerol and 1 mM TCEP on an ÄKTA micro system (Cytiva). All samples were eluted under isocratic conditions at 4°C in SEC buffer (20 mM HEPES pH 6.8, 300 mM NaCl, 5 % (v/v) glycerol and 1 mM TCEP) at a flow rate of 0.15 ml/min. Elution of proteins was monitored at 280 nm, 555 nm and 647 nm in case of ^mScarlet^CENP-TW and CENP-SX^Alexa647^. 100 µl fractions were collected and analyzed by SDS-PAGE and Coomassie blue staining. In experiments where fluorescent labeled proteins were used, the in-gel detection of the fluorescence was detected using a BioRAD chemiDoc MP Imaging System (BioRAD, Hercules, US-CA). To detect the formation of a complex, proteins were mixed at the concentrations of 5 µM in 50 µl, incubated for at least 1 hour on ice, subjected to SEC then analyzed by SDS-PAGE.

### Amylose-resin pull-down assay

The proteins were diluted with binding buffer [20 mM HEPES pH 6.8, 300 mM NaCl, 2.5% glycerol, 1 mM TCEP, and 0.01% Tween] to 3 µM concentration in a total volume of 50 µl and mixed with 20 μl of amylose beads (New England Biolabs, Ipswich, US-MA). After mixing the proteins and the beads, 20 µl were taken as input. The rest of the solution was incubated at 4°C for 1 hour on a thermomixer (Eppendorf, Hamburg, Germany) set to 1000 rpm. To separate the proteins bound to the amylose beads from the unbound proteins, the samples were centrifuged at 800g for 3 min at 4°C. The supernatant was removed, and the beads were washed four times with 500 µl of binding buffer. After the last washing step, 20 µl of 2× SDS-PAGE sample loading buffer was added to the dry beads. The samples were boiled for 5 min at 95°C and analyzed by SDS-PAGE and Coomassie staining. Gel densitometry was carried out with Image Lab (BioRAD, Hercules, US-CA).

### In vitro assembly of CCAN

Reconstitution of human recombinant CCAN particles was performed as previously published in Pesenti et al., 2018. In brief, a stoichiometric amount of purified CENP-LN, CENP-CHIKM, CENP-OPQUR and CENP-TWSX complexes were incubated at around 15 µM at 4°C for minimum 1 hour and purified by SEC using a buffer containing 20 mM HEPES pH 6.8, 300 mM NaCl, 2.5 % (v/v) glycerol and 1 mM TCEP.

### Sample preparation for electron microscopy

The reconstituted CCAN particles were stabilized via the GraFix method (Kastner et al., 2008). Two 4-ml gradients ranging from 20 to 50 % glycerol in 20 mM HEPES pH 6.8, 300 mM NaCl and 1mM TCEP were set up. In one of the gradients, the 50 % solution contained 0.125% of glutaraldehyde. Around 100 µL of sample at 15 µM was applied to each gradient and centrifuged by ultracentrifugation at 45 000 rpm at 4°C in SW 60 Ti Swinging-bucket rotor (Beckman Coulter, Palo Alto, US-CA) for 16 hours. The samples from both gradients were fractionated in 150 μL fractions, and cross-linker containing fractions were quenched by addition of 100 mM Tris pH 6.8. All fractions were analyzed by SDS-PAGE and Coomassie blue staining. The fractions of interest were dialyzed two times against 2 l of 20 mM HEPES pH 6.8, 300 mM NaCl, and 1mM TCEP buffer for 16 hours and 2 hours, and concentrated to around 2 mg/ml using Amicon Ultra 0.5 ml-100 kDa cutoff (Millipore, Burlington, US-MA).

### Negative stain electron microscopy

Negative stain specimens were prepared as described previously (Brocker et al., 2012): The cross-linked CCAN samples were diluted in 20 mM HEPES pH 6.8, 300 mM NaCl, and 1mM TCEP buffer to adjust the particle density. 4 µL of the sample were absorbed at 25 °C for 1 min onto freshly glow-discharged 400 mesh carbon-coated copper grids (G2400C, Plano GmbH, Wetzlar, Germany). Excess sample was blotted by touching a Whatman filter paper and washed with three droplets of water and exposed to freshly prepared 0.75 % uranyl formate solution (SPI Supplies/Structure Probe, West Chester, PA) for about 1 min. Excess negative stain solution was blotted and the specimen air-dried. Specimens were inspected with a JEM1400 microscope (Jeol, Tokio, Japan) equipped with a LaB_6_ cathode and operated at an acceleration voltage of 120 kV. Digital micrographs were recorded using a 4k x 4k CMOS camera F416 (TVIPS, Gauting, Germany).

### Cryo-EM grid preparation and data acquisition

Grids were prepared using a Vitrobot Mark IV (Thermo Fisher Scientific) at 13 °C and 100 % humidity. 4 µl of CENP-16 supplemented with 0.0025 % Triton were applied to glow-discharged UltrAuFoil R1.2/1.3 grids and excess liquid removed by blotting (3.5 seconds at blot force -3) before vitrification in liquid ethane. For dataset I, CENP-16 was used at a concentration of 1.5 mg/ml. For dataset II, CENP-14 (without CENP-SX) was used at 0.9 mg/ml. The CCAN sample used for dataset II also contained a 145-bp DNA and CENP-A:H4. Elongated DNA was visible in the micrographs, but no density for DNA or CENP-A:H4 was identifiable in the 2D classes. Dataset I was acquired on a Titan Krios electron microscope (Thermo Fisher Scientific) equipped with a field emission gun. For this first dataset, 1540 movies were recorded on a K3 camera (Gatan) operated in super-resolution mode at a nominal magnification of 130000, resulting in a super-resolution pixel size of 0.35 Å. A Bioquantum post-column energy filter (Gatan) was used for zero-loss filtration with an energy width of 20 eV. Total electron exposure of 76.8 e^-^/Å^2^ was distributed over 80 frames. Data were collected using the automated data collection software EPU (Thermo Fisher Scientific), with two exposures per hole and a set defocus range of -0.6 to -1.2 µm. The second dataset was recorded on a Cs-corrected Titan Krios microscope equipped with a K3 camera (Gatan) and a Bioquantum post-column energy filter with a slit width of 14 eV operated in super-resolution mode at a nominal magnification of 105000, corresponding to a super-resolution pixel size of 0.34 Å. A total exposure of 55.8e^-^/Å^2^ was distributed over 60 frames. 2678 movies were collected using EPU (Thermo Fisher Scientific), with two exposures per hole and a set defocus range of −0.6 to −1.2 µm. For both datasets, phase contrast was induced by using a volta phase plate in the back focal plane. Details of data acquisition parameters can be found in Table S1.

### Cryo-EM data processing

On-the-fly data pre-processing, including correction of beam-induced motion and dose-weighting by MotionCor2 (Zheng et al., 2017), CTF parameter estimation using CTFFIND4 in movie mode (Rohou and Grigorieff, 2015), and particle picking using a custom neural network in SPHIRE-crYOLO (Wagner et al., 2019), was performed within TranSPHIRE (Stabrin et al., 2020). For the high-resolution dataset (dataset I in Figure 1 – Supplement 3), template-free particle picking by crYOLO (Wagner et al., 2019) in the 1540 micrographs greatly improved after re-training with 1354 manually picked particles, resulting in 140910 particle coordinates. 2-fold binned particles were extracted in SPHIRE (Moriya et al., 2017) using a box size of 220×220 pixels. 2D classification was performed in ISAC with a class size limit of 500 particles, a particle radius of 105 pixels and using the VPP option. 45 beautified 2D class averages which had been filtered to 8 Å were used to generate an initial 3D model in RVIPER. In parallel, the dataset was processed in RELION 3.1.2 (Fernandez-Leiro and Scheres, 2017; Nakane et al., 2018) using a box size of 384×384 pixels for extraction and 200 classes for 2D classification and initial model generation. The 95522 selected particles and the resulting initial model could be refined both in MERIDIEN and RELION, yielding a 7 Å reconstruction. The centered particles were then re-extracted without binning and using a box size of 384 pixels, and all further processing steps were performed in RELION. 3D classification with four classes yielded one class with 25206 particles which showed high-resolution features in the center of the particle. The quality of the reconstruction was improved by Bayesian polishing, resulting in an increased global resolution of 5 Å.

This reconstruction was further improved by multi-body refinement in RELION using two masks covering the majority of the HIKMLN or OPQUR subcomplexes, i.e. omitting the more flexible QU- and HIK-TW-“heads”, resulting in focused reconstructions with resolutions of 4.6 Å for HIKMLN and 6.9 Å for OPQUR. Segmenting the volumes further (including e.g. additional maps for the “heads”) did not improve the quality of the reconstructions. The multi-body refinement was especially important for improvement of the resolution of the OPQUR part. As evident from local resolution estimation with RELION, the quality of the reconstruction varies greatly between the well-ordered CENP-M/L/N interface with local resolution of 3.7 Å, and more peripheral parts which are more flexible and less well resolved. The angular distribution showed that the particles had a moderate fraction of preferred orientations along the shortest axis of the particles (Figure 1 – Supplement 4). DeepEMhancer (Sanchez-Garcia et al., 2021) was used to further enhance the maps for model building.

The second dataset (dataset II in Figure 1 – Supplement 3) had lower resolution although the initial 2678 micrographs yielded more particles compared to the first dataset (233598 vs 140910) when picked with the re-trained crYOLO model. DNA strands were visible in the micrographs, but no nucleosomes (no H2A:H2B was present). For extraction, a larger box size of 512 pixels (corresponding to 358.4Å) was chosen to potentially detect more different conformations compared to the first dataset. Particles were subjected to 2D classification in RELION, and an initial model was calculated, also in RELION, using 117473 particles assigned to good 2D classes. Subsequent 3D classification into 4 classes yielded one class (with 44216 assigned particles) suitable for 3D refinement. The quality of the refined model could not be improved by further 2D classification, Bayesian polishing, or CTF refinement. Although the overall resolution was lower, this dataset showed much clearer density for the HIK and QU heads, including a tentative density for the CENP-TW complex that fits very well to the position of TW in the X-ray structure of the HIK-TW complex (PDB ID 6WUC). This assignment is corroborated by Alphafold2 predictions that indicate a strong interaction between the HIK head and TW as compared to the CENPA/H4 dimer (I.R.V, unpublished results). RELION multibody refinement with three masks covering HIKTW, MLN, and OPQUR, respectively, resulted in resolutions of 10.2Å, 10.2Å and 10.6Å for the three maps. Local resolution estimated with RELION ranged between 8-25 Å (Figure 1 – Supplement 4). The angular distribution showed no pronounced preferred orientations. Since crYOLO picked practically all particles and the 2D classes did not show any evidence for the presence of DNA or nucleosomes, any stable association of DNA or CENP-A/H4 with the CCAN can be excluded.

### Model building and structure refinement

Crystal structures of human CENP-M (PDB ID 4WAU), human CENP-N (PDB ID 6EQT) as well as *ab initio* models for human CENP-H, K, I, L, N, O, P, Q, U, R, T, and W predicted by the Tencent tFold server (https://drug.ai.tencent.com) were initially docked as rigid bodies into the reconstruction of dataset I sharpened by DeepEMhancer and then locally adjusted using Coot (Emsley et al., 2010) and Chimera (Yang et al., 2012). The cryo-EM structures of the yeast CCAN (PDB IDs 6QLD and 6QLE) and the yeast CTF3 complex with CNN1-WIP1 (PDB ID 6WUC) were also used to guide the modeling. Subsequently, *ab initio* models predicted using Alphafold2 were used to further improve the structure model. These Alphafold models (“Colabfold”, AF2, and AF2-multimer (Evans et al., 2021; Jumper et al., 2021; Mirdita et al., 2021) greatly facilitated the sequence assignment in regions with lower local resolution. Density for a short helix at the interface of CENP-I and CENP-K was built as a polyalanine model since the sequence could not be assigned reliably. Possible assignments include CENP-C, the N-terminus of CENP-H, CENP-P or CENP-O, the latter slightly more likely since AF2-multimer predicted this interaction with high confidence. In contrast, the extension of the CENP-N *β*-sheet by the CENP-C region 303-FIID-306 was reliably predicted by AF2-multimer and allowed unambiguous interpretation of the corresponding electron density for CENP-C. The position of the CENP-TW complex was derived by superposing an AF2-prediction of the CENP-HIK “head” in complex with CENP-TW (which had exactly the same arrangement as the model of the yeast CTF3-CNN1-WIP1 complex (PDB ID 6WUC)) and subsequently docking it into the reconstruction of dataset II. The final model was optimized by geometry minimization and real-space refinement with PHENIX (Adams et al., 2010) and evaluated with COOT, PHENIX and MolProbity (Chen et al., 2010). Figures were prepared using Chimera and PyMOL (Molecular Graphics System, 2.0.3, Schrodinger). Structural sequence alignments were performed with Chimera.

### Cell culture and immunofluorence

HeLa Flip-In T-REx EGFP-CENP-M (Basilico et al., 2014) were maintained in DMEM with 10% tetracycline-free FBS (Pan Biotech), supplemented 50 μg/ml Penicillin/Streptomycin (PAN Biotech), and 2 mM L-glutamine (PAN Biotech). For immunofluorence experiments, cells were grown on coverslips pre-coated with 0.01% poly-D-lysine (Sigma-Aldrich). Exogenous gene expression was induced by adding 50 ng/ml doxycycline (Sigma, St. Louis, Missouri, United States) to the media for 24h before fixation. Cells were fixed with ice-cold Methanol for 1 minute, then washed 3 times for 5 minutes with 1 X PBS + 0.1% Tween-20 (PBST). Cells were blocked for 20 minutes at room temperature in PBST + 5% BSA, and then were incubated in primary antibodies diluted in PBST + 1% BSA overnight at 4C. The following morning, coverslips were washed 3 times for 5 minutes with PBST and then incubated for 30 minutes at room temperature with secondary antibodies diluted in PBST + 1% BSA. Finally, coverslips were washed 3 times for 5 minutes in PBST and quickly rinsed in distilled water before mounting. The following primary antibodies were used: anti-CyclinB1 (rabbit monoclonal antibody, Abcam, #ab32053, 1:1000), CREST/anti-centromere antibody (human autoimmune serum, Antibodies Inc., #15-234, 1:1000), anti-PCNA (mouse monoclonal antibody, Cell Signalling, #2586S, 1:1000), anti-Tubulin (rat monoclonal antibody, Abcam, #6160, 1:500). The following secondary antibodies were used: donkey anti-human DyLight 405 (Jackson ImmunoResearch, 1:000), donkey anti-mouse Alexa Fluor® 647 (Invitrogen, 1:1000), donkey anti-rabbit Rhodamine Red (Jackson ImmunoResearch, 1:000), donkey anti-rat Rhodamine Red (Jackson ImmunoResearch, 1:000), GFP-Booster Alexa Fluor® 488 (Chromotek, gb2AF488-50, 1:1000), goat anti-human Alexa Fluor® 647 (Jackson ImmunoResearch, 1:000). DNA was stained with 0.5 μg/ml DAPI (Serva) and Mowiol (Calbiochem) was used as mounting media.

### Accession codes

The coordinates and map of the high- and low-resolution CCAN models have been submitted to the Protein data bank with ID 7QOO and to the EMDB with IDs EMD-14098 for the high-resolution dataset and EMD-14099 for the low-resolution dataset.

## Acknowledgements

We are grateful to Oliver Hofnagel for help with data collection, to Julia Schweighofer and Sabine Wohlgemuth for help with the production of CENP-LN and CENP-TW, and to Karolin Luger, Keda Zhou, and the Musacchio and Raunser laboratories for helpful discussions and sharing of unpublished results. This work was supported by the Max Planck Society (to A.M. and S.R.). D.C. acknowledges funding by the European Molecular Biology Organization (EMBO) through an EMBO Long-Term Fellowship (ALTF 439-2019). A.M. acknowledges funding by the Marie-Curie Training Network DivIDE (project number 675737), European Research Council (ERC) through Synergy Grant 951439 (BIOMECANET) and formerly AdG 669686 (RECEPIANCE). The authors declare no competing financial interests.

## Author contributions (following CRediT model)

**Conceptualization:** Andrea Musacchio

**Data curation:** N/A

**Investigation**

**Sample preparation:** Marion E. Pesenti
**Data collection and processing:** Daniel Prumbaum, Tobias Raisch, Ingrid R. Vetter
**Structure determination and model building:** Ingrid R. Vetter
**Cell Biology:** Duccio Conti

**Funding acquisition:** Andrea Musacchio, Stefan Raunser

**Project Administration:** Andrea Musacchio, Stefan Raunser

**Resources:** Ingrid Hofmann, Doro Vogt

**Supervision:** Andrea Musacchio, Stefan Raunser, Ingrid R. Vetter

**Validation:** Andrea Musacchio, Stefan Raunser, Ingrid R. Vetter

**Visualization:** Duccio Conti, Marion E. Pesenti, Tobias Raisch, Ingrid R. Vetter

**Writing – original draft:** Andrea Musacchio

**Writing – review & editing:** All authors

## Figure Supplement Legends

**Figure 1 – Supplement 1.**
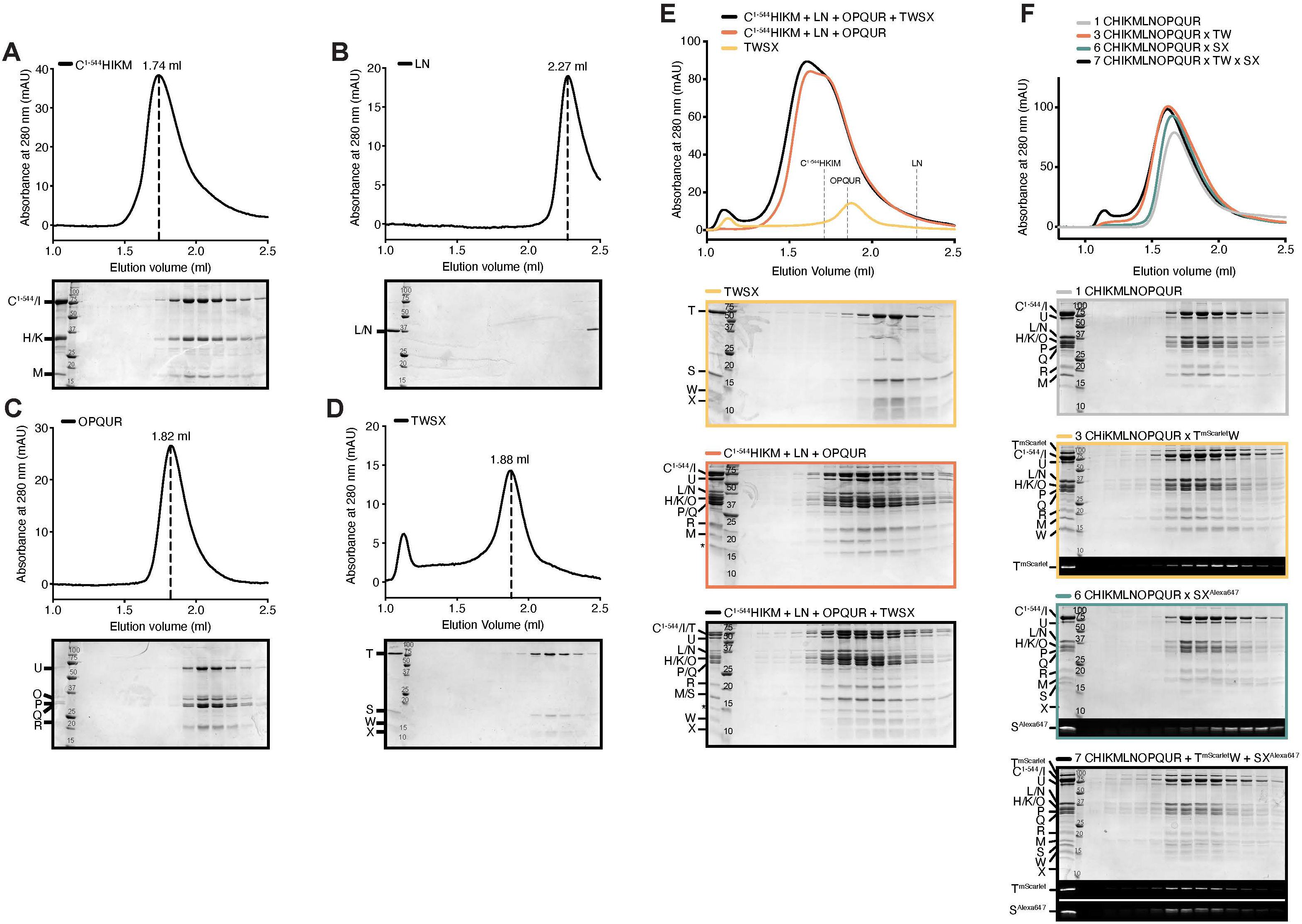
Biochemical reconstitution of human CCAN. Size-exclusion chromatography on a Superose 6 5/150 column of the indicated CCAN complexes and subcomplexes **A**) CENP-C^1-544^HIKM. **B**) CENP-LN. **C**) CENP-OPQUR. **D**) CENP-TWSX. DNA binding is demonstrated by the modest left shift when samples was incubated with DNA. **D**) CENP-OPQUR. DNA binding is demonstrated by a left shift when sample was incubated with DNA. **E**) Comparison of elution profiles of CENP-TWSX, CENP-12, and CENP-16. **F**) CENP-TW binds CENP-12, but CENP-SX requires CENP-TW to interact with it.

**Figure 1 – Supplement 2.**
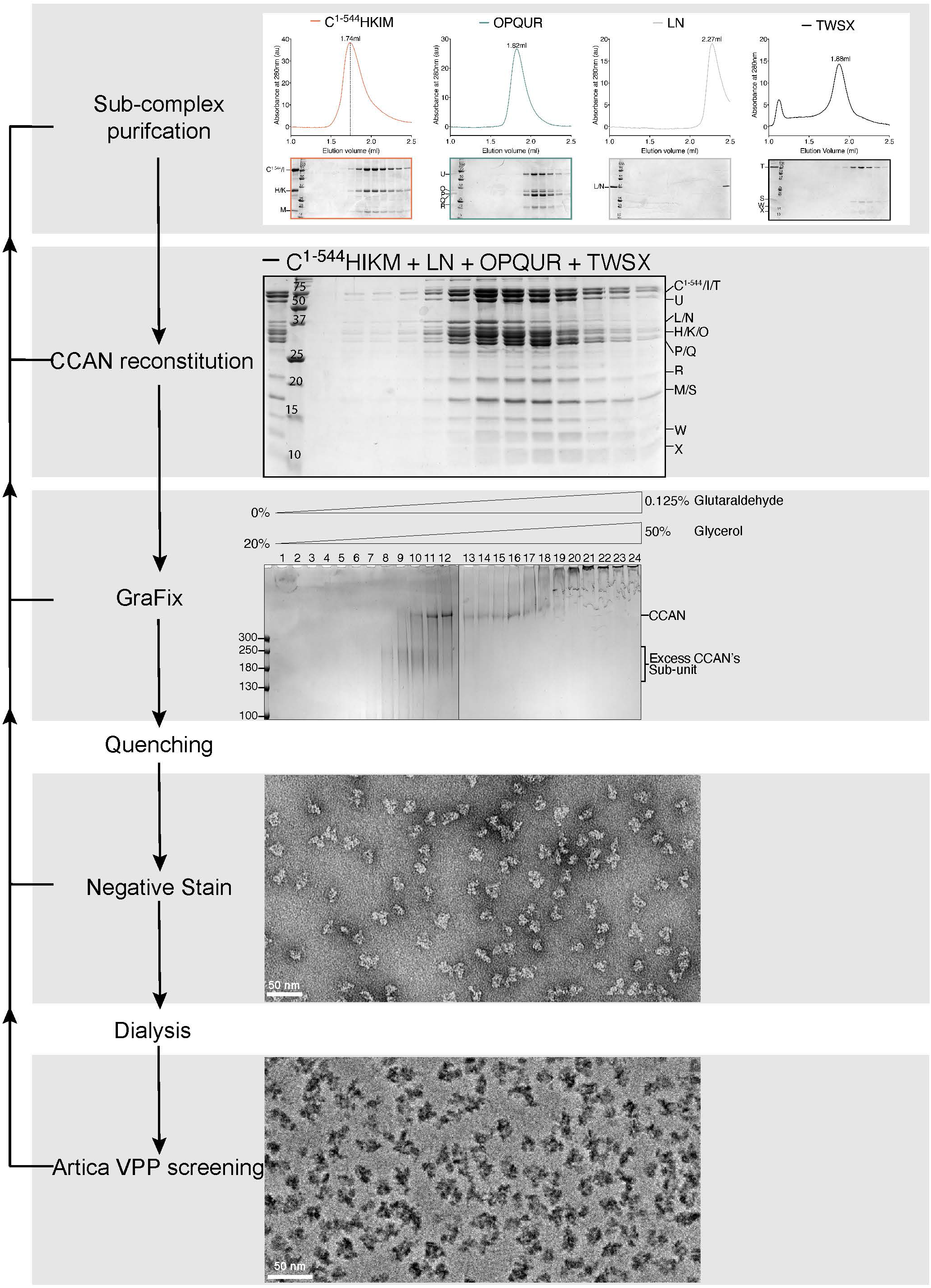
Pipeline of sample preparation. The figure presents an outline of the various steps of sample preparation preceding high-resolution cryo-EM data collection.

**Figure 1 – Supplement 3.**
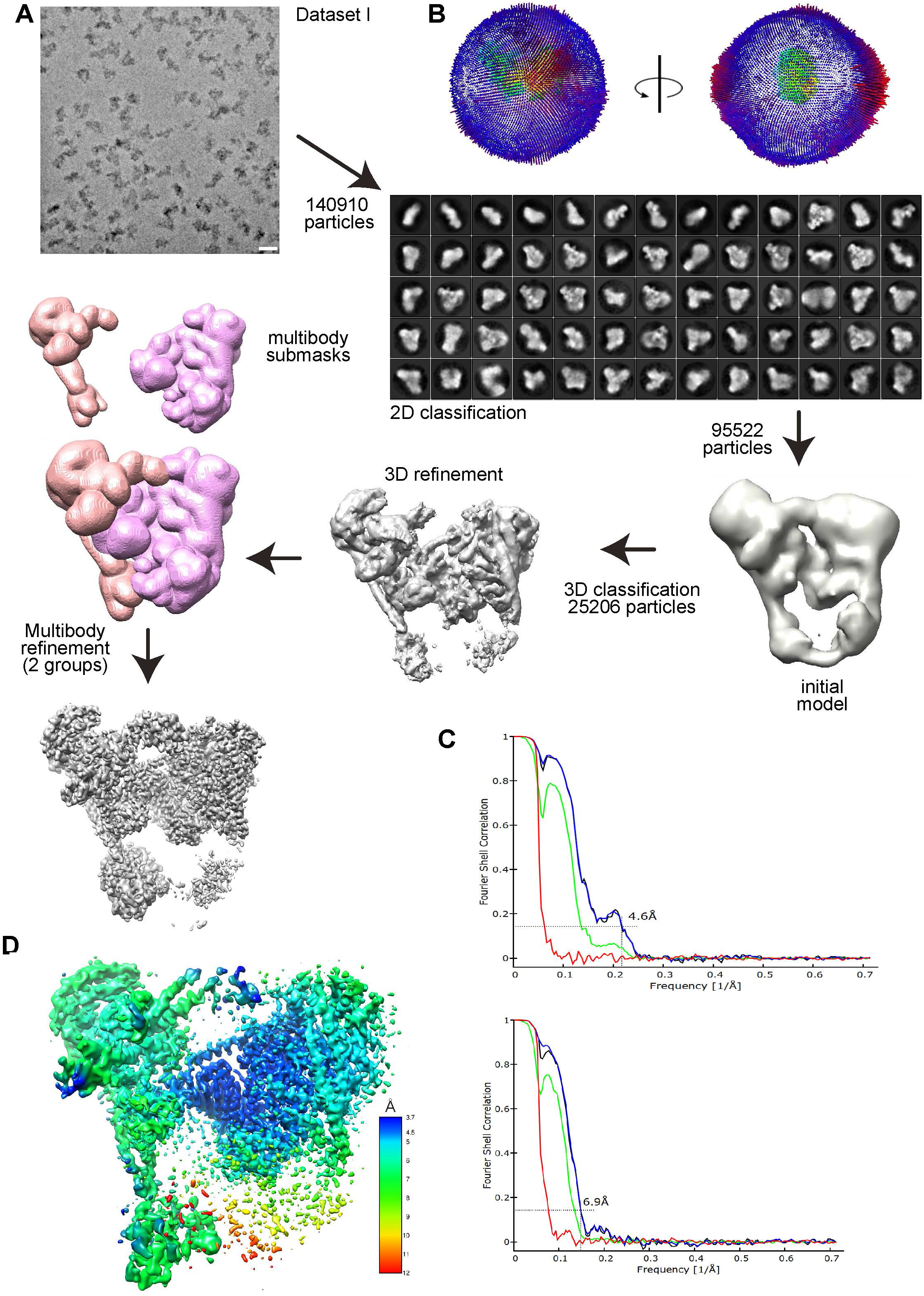
Data processing flowchart for Dataset I. **A**) Processing flowchart for the high-resolution dataset I including an exemplary micrograph (scale bar = 20 nm) and a subset of selected 2D classes of CENP-16. The last (grey) map shows the final reconstruction and was obtained by combining the two focused maps from multibody refinement (using the ‘vop maximum’ command in UCSF Chimera). **B**) Angular distribution of the particles shown in two positions rotated 90° to each other. **C**) Fourier shell correlation (FSC) plots between two independent half-maps for each of the two bodies used in the multibody refinement procedure, according to the FSC=0.143 criterion. The dashed line indicates the 0.143 FSC criterion. red: phase-randomized map, green: unmasked map, blue: masked map, black: corrected map. **D**) Local resolution estimates by RELION for dataset I plotted on the two multibody reconstructions in a rainbow-colored gradient from blue (3.7 Å) to red (12 Å). The map differs from the final reconstruction depicted in panel A since the latter shows the combined sub-maps, further modified by DeepEMhancer.

**Figure 1 – Supplement 4.**
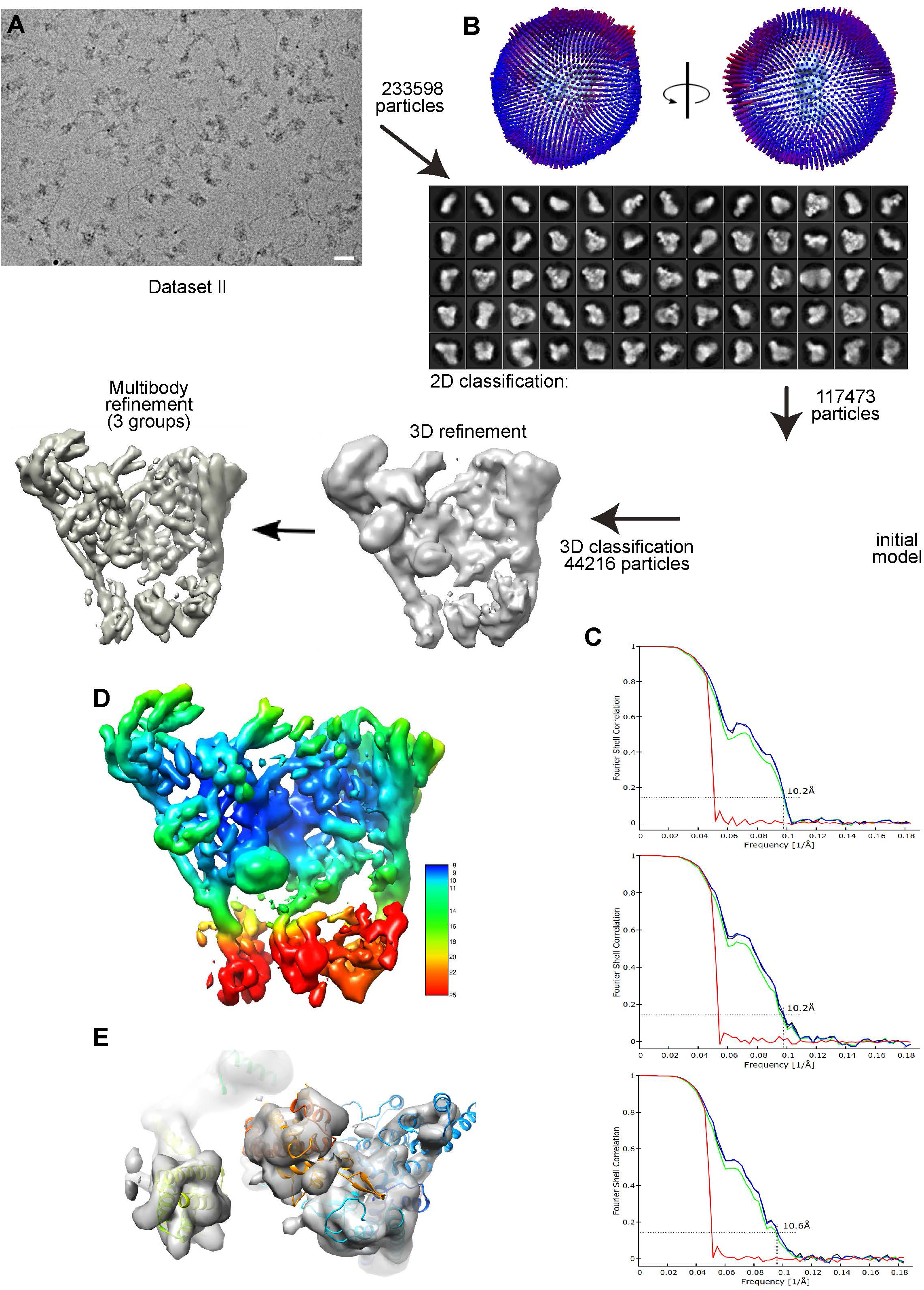
Data processing flowchart for Dataset II. **A**) Processing flowchart for the low-resolution dataset II including an exemplary micrograph and a subset of selected 2D classes of CENP-14. **B**) Angular distribution of the particles shown in two positions rotated 90° to each other. **C**) Fourier shell correlation (FSC) plots between two independent half-maps for each of the three bodies used in the last multibody-refinement, indicating the resolution of the three groups in the multibody refinement according to the FSC=0.143 criterion. The dashed line indicates the 0.143 FSC criterion. red: phase-randomized map, green: unmasked map, blue: masked map, black: corrected map. **D**) Local resolution estimates by RELION plotted on the multibody reconstructions in a rainbow-colored gradient from blue (8 Å) to red (25 Å). **E**) Representative density of CENP-QU head (left), the CENP-TW complex (middle) and the HIK head (right), as seen from the “bottom” towards the top of the CCAN complex in D).

**Figure 1 – Supplement 5.**
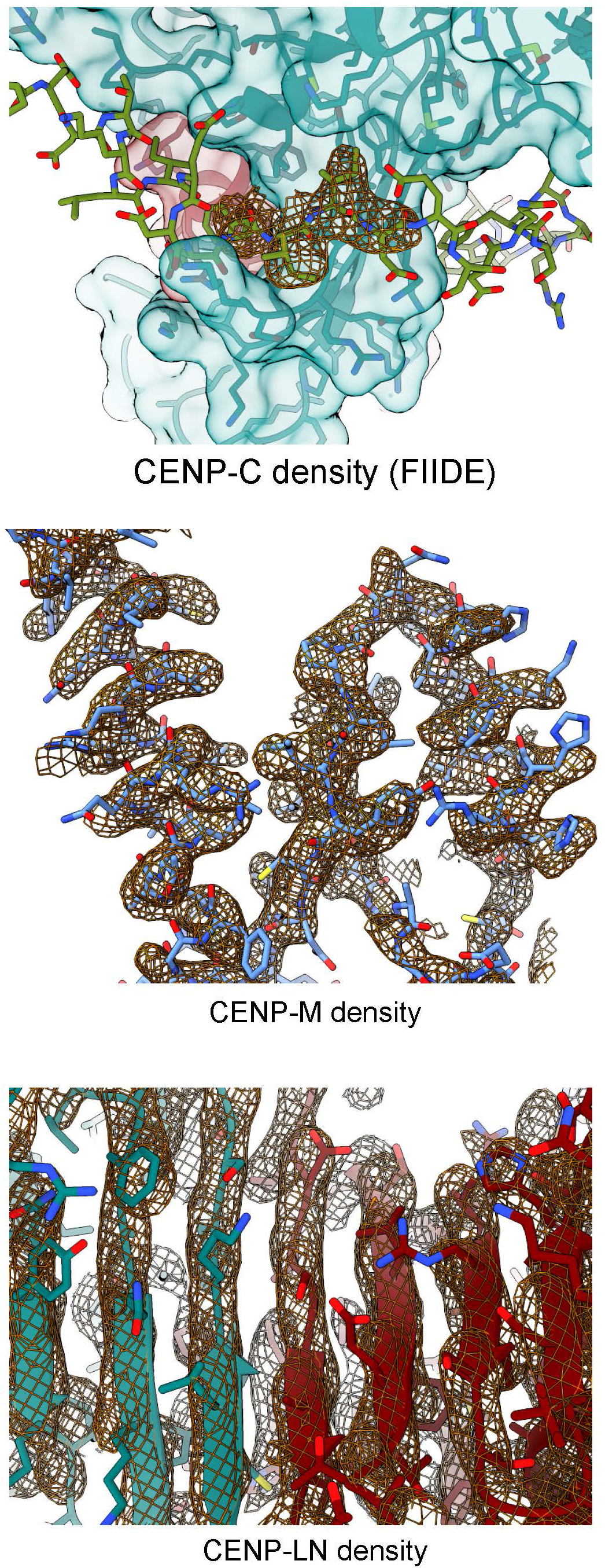
Close-ups of density maps. **A**) Density for CENP-C 303-FIID-306 bound to CENP-N. **B**) Representative density of the region with highest resolution around CENP-M. **C**) Density at the CENP-LN β-sheet interface.

**Figure 1 - Supplement 6.**
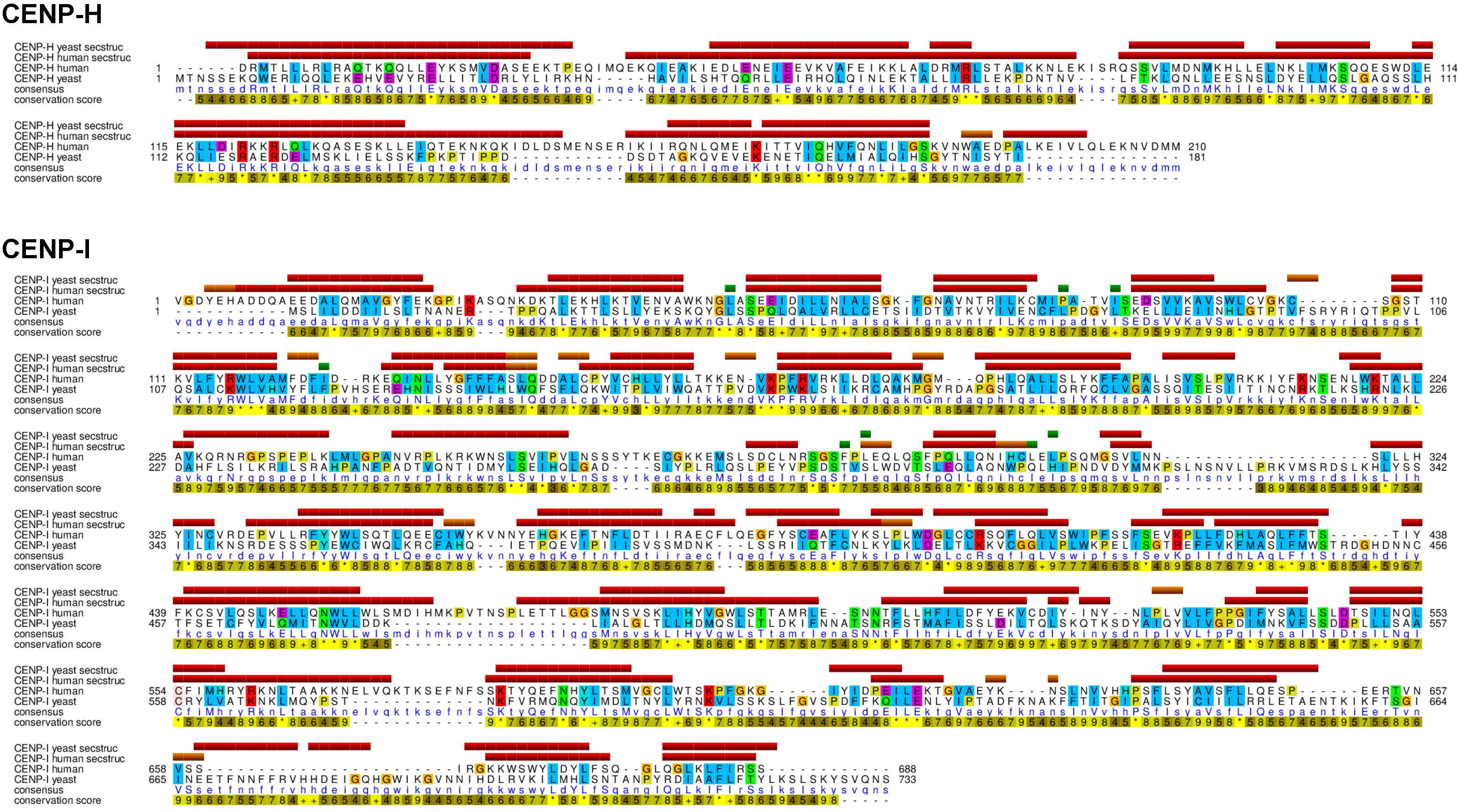
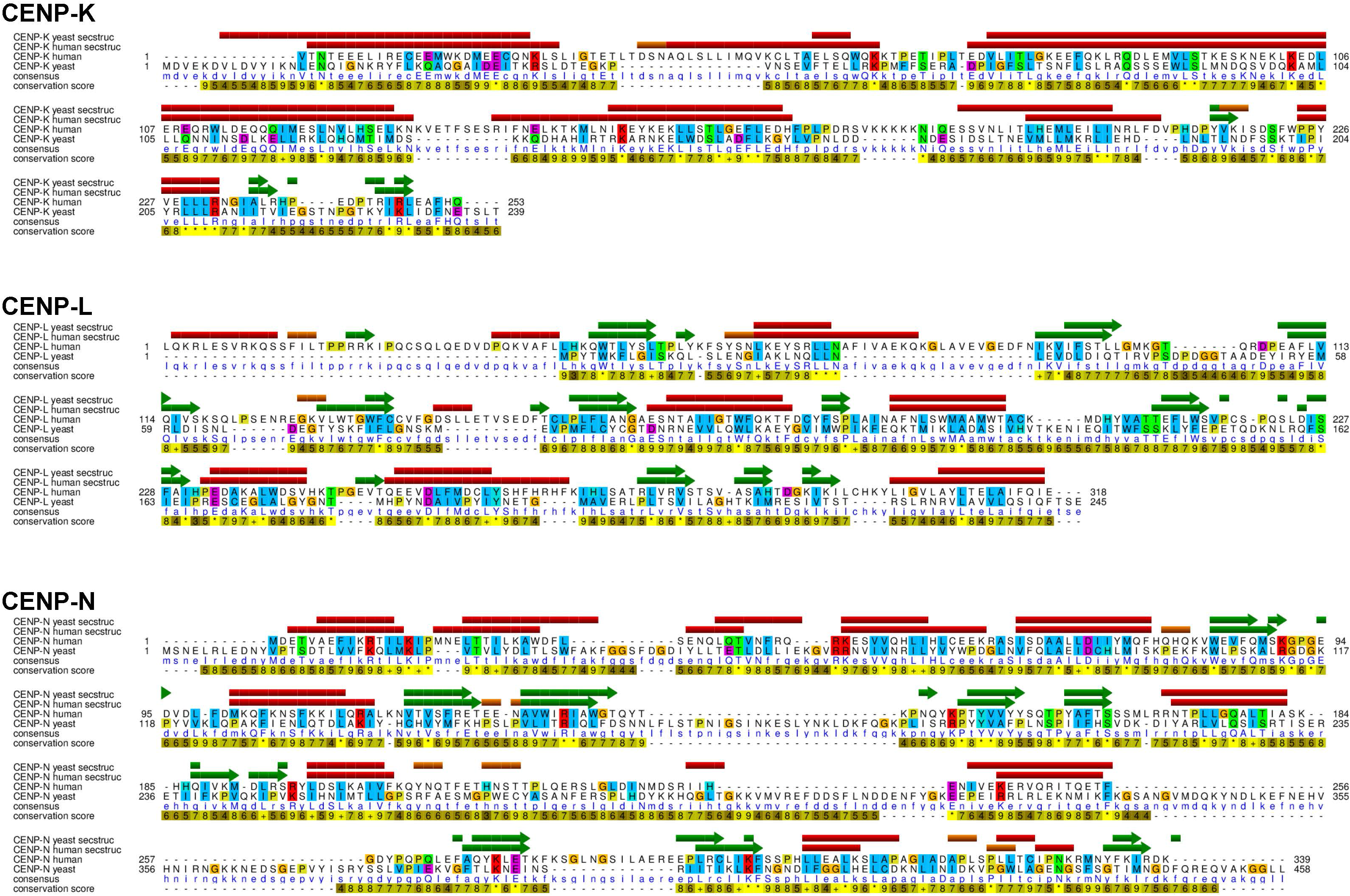
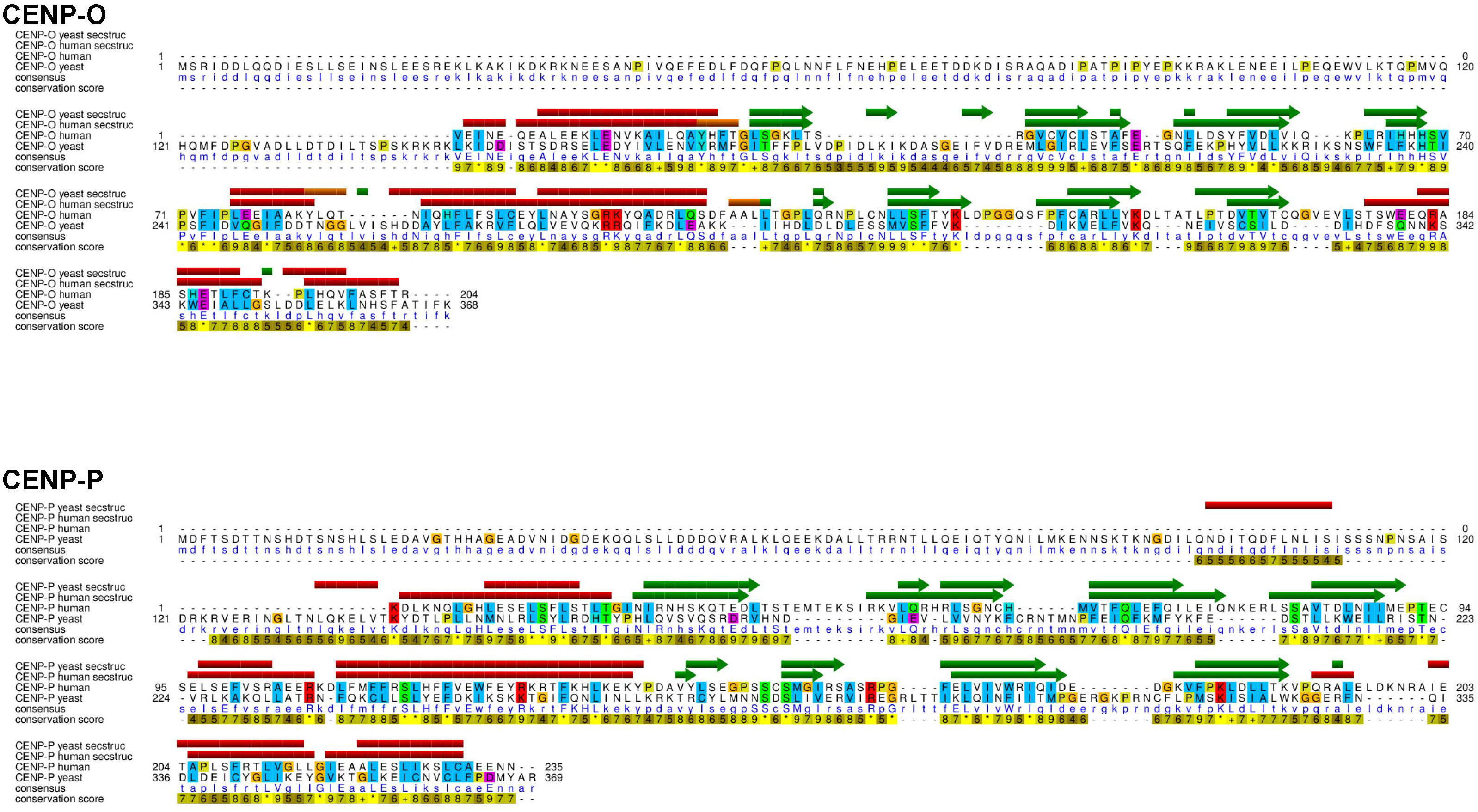
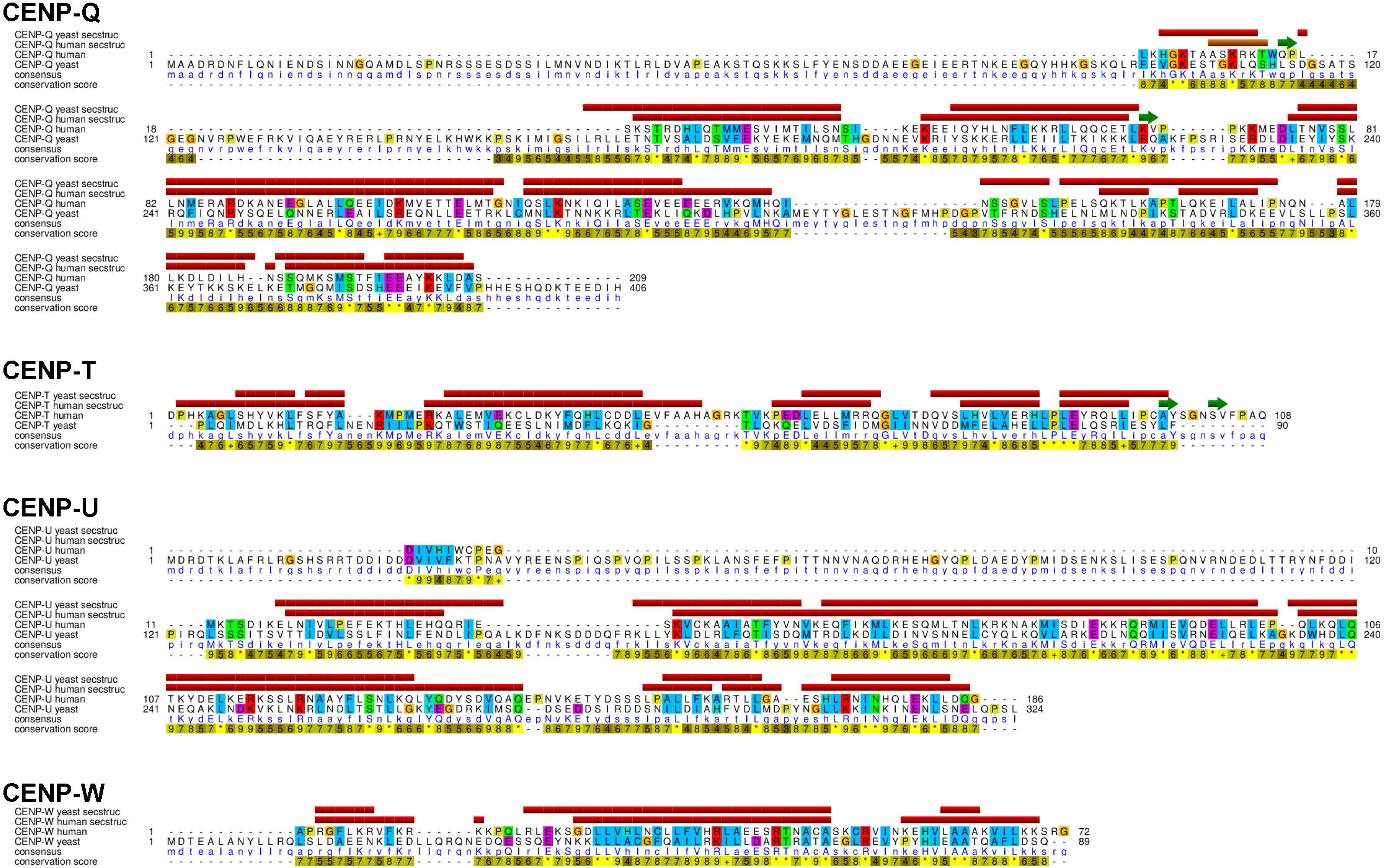
Sequence alignments of human and yeast CCAN. Secondary structure elements of yeast CCAN (PDB ID 6QLE) were aligned to the human CCAN structure by keeping the molecular shapes and relative orientations as similar as possible. Structure-based alignments were computed by Chimera. The secondary structures for the yeast proteins were assigned by DSSP (Kabsch and Sander, 1983) from the original yeast structures with PDB ID 6QLE and 6WUC. *α*-helices are shown in red, 3_10_ helices in orange, and *β*-sheets in green. Sequences are colored according to the ClustalX coloring scheme.

**Figure 2 – Supplement 1.**
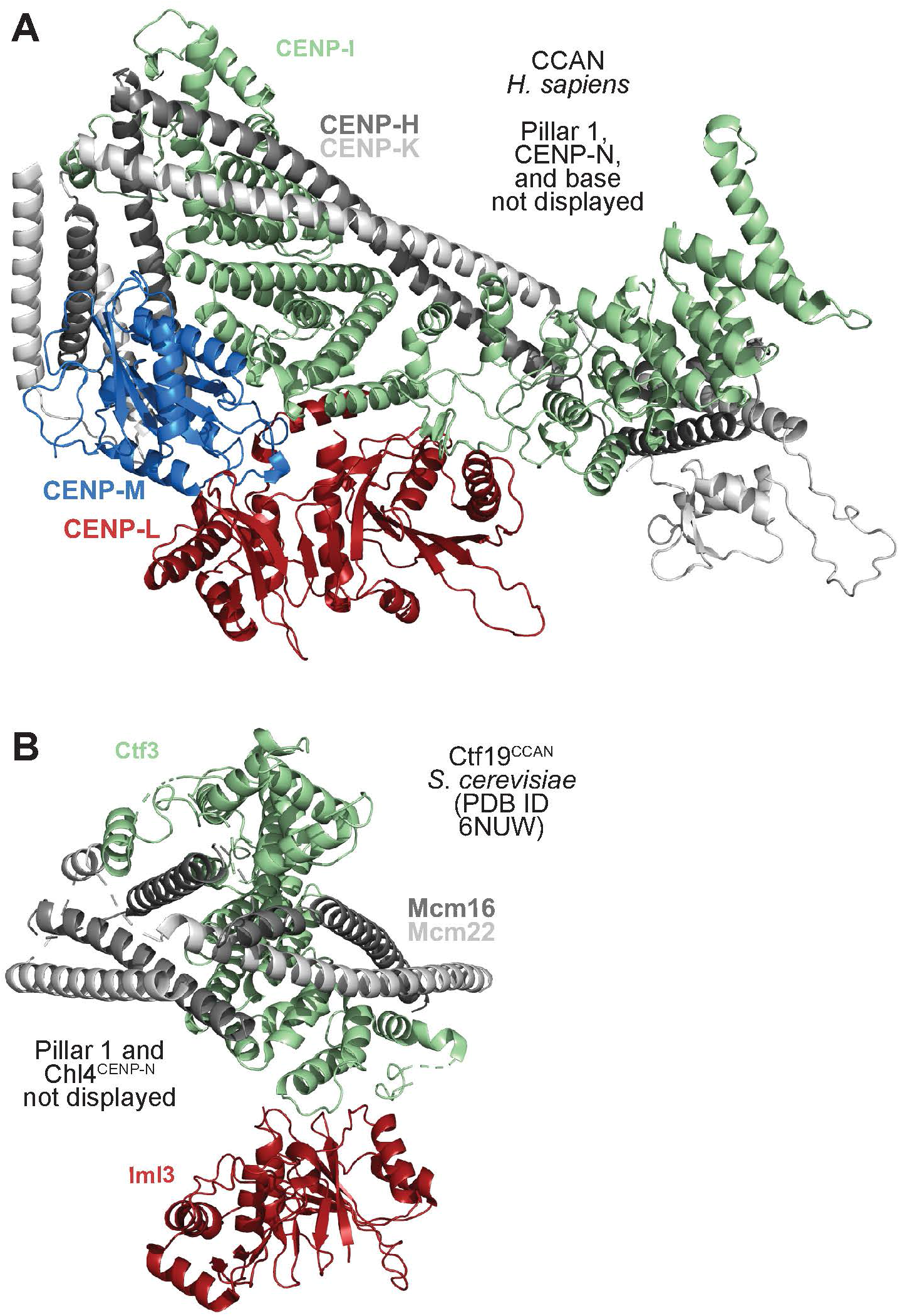
Comparative interaction of pillar 2 with the vault. **A**) Cartoon model of pillar 1 and CENP-L in the vault for human CCAN demonstrates extensive interactions, as already illustrated in Figure 2E-F. **B**) The equivalent region in the *S. cerevisiae* complex shows a much more modest interface that is limited to the Iml3^CENP-L^:Ctf3^CENP-I^ pair and that further emphasizes the role of CENP-M as stabilizer of the human complex.

**Figure 2 – Supplement 2.**
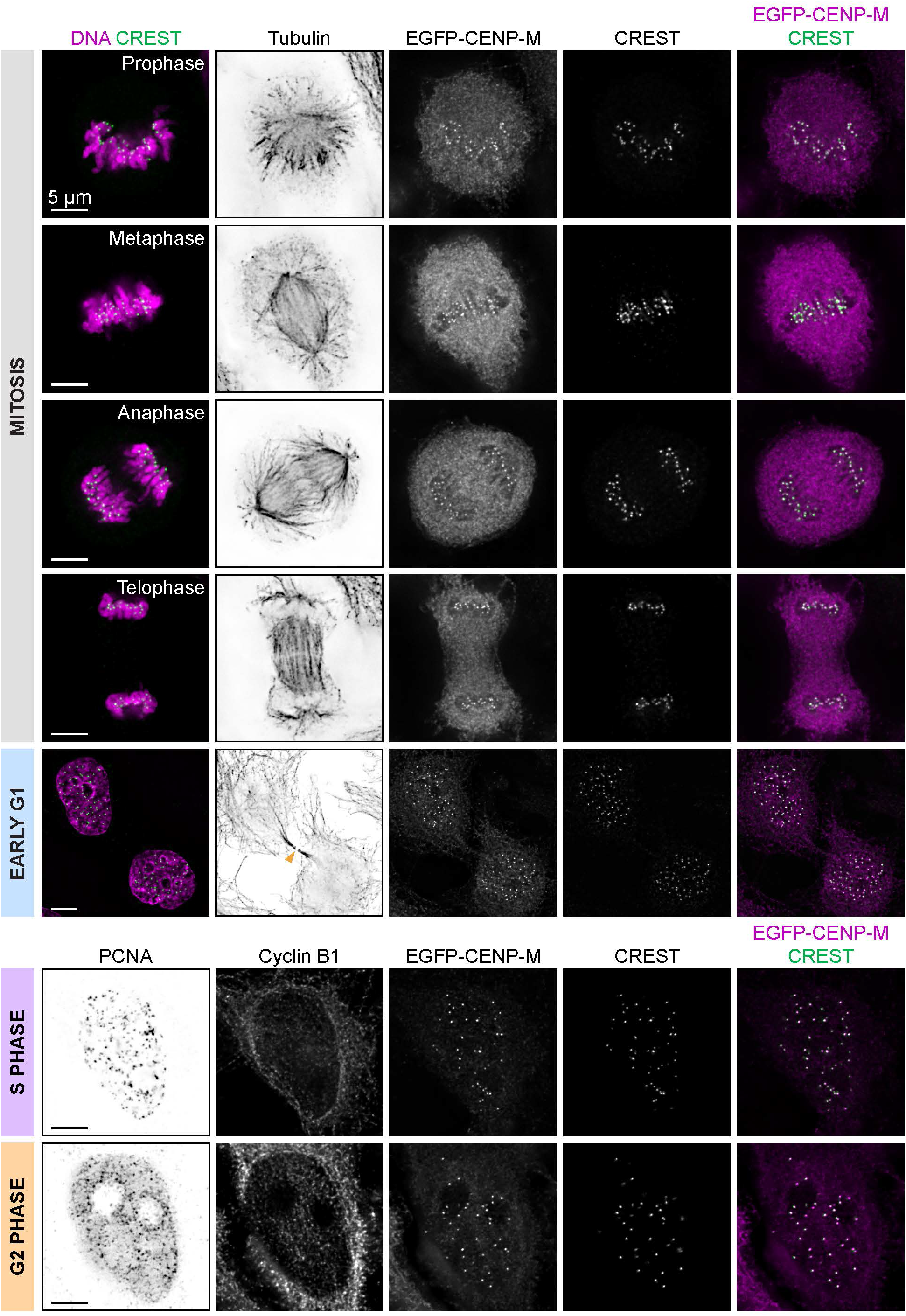
Gallery of EGFP-CENP-M cell cycle localization. Maximum intensity projections of 5 x 0.2 µm Z-stacks. For Early G1 cells, the orange arrowhead indicates the spindle midbody remnant. Tubulin and DNA were the chosen markers to illustrate mitotic figures. PCNA and cytosolic Cyclin B1 were the chosen markers to illustrate S-phase and G2 phase.

**Figure 3 – Supplement 1.**
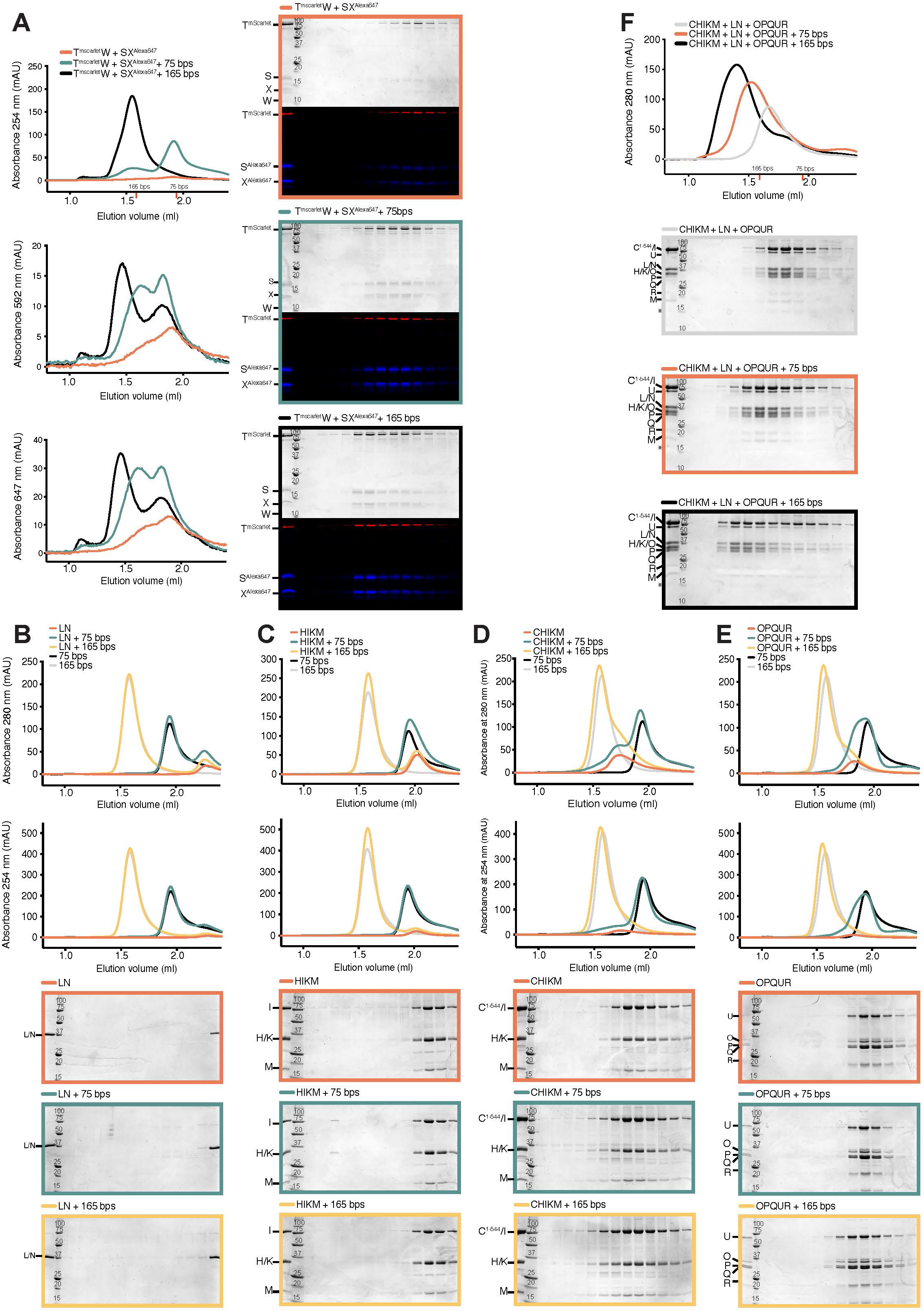
DNA binding by CCAN subunits. Size-exclusion chromatography on a Superose 6 5/150 column of the indicated samples to monitor protein:DNA binding using 75-bps and 165-bps DNA. **A**) CENP-LN. **B**) CENP-HIKM. **C**) CENP-C^1-544^HIKM. DNA binding is demonstrated by the modest left shift when samples was incubated with DNA. **D**) CENP-OPQUR. DNA binding is demonstrated by a left shift when sample was incubated with DNA. **E**) CENP-TWSX bound strongly to DNA. **F**) The combined CCAN subunits in CENP-12 bound DNA with substantial affinity.

**Figure 4 – Supplement 1.**
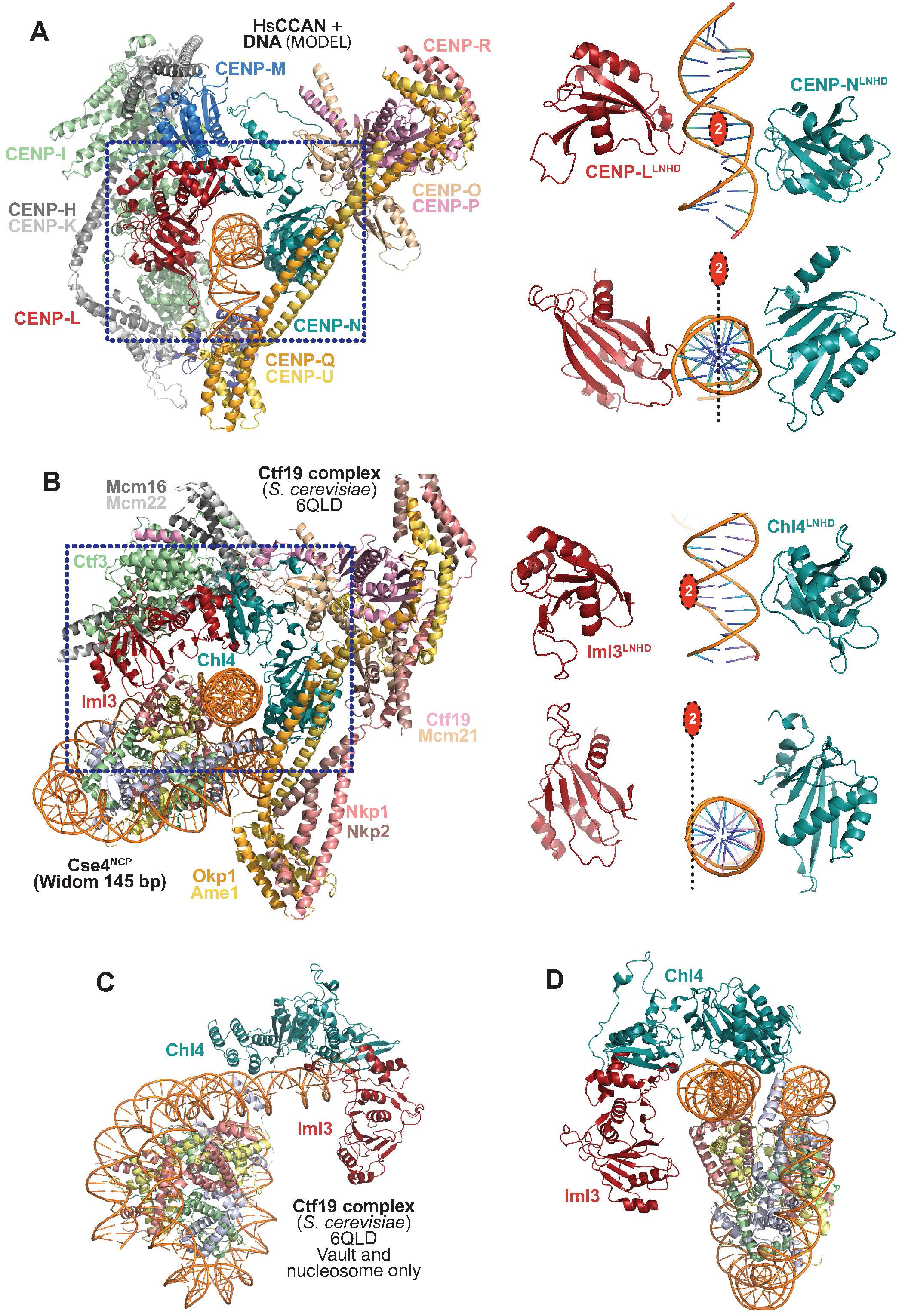
2-fold pseudosymmetry in the human vault. **A**) *Left*: Cartoon model of human CCAN bound to DNA. The DNA fragment was extracted from PDB ID 6C0W (CENP-N N-terminal region bound to CENP-A nucleosome (Pentakota et al., 2017)) from the DNA gyre facing CENP-N after superposition of CENP-N. *Right*: A 2-fold pseudosymmetry axis links the two LNHDs and the modelled DNA. Related views are shown in Figure 4F-G. **B**) *Left*: View of the *S. cerevisiae*’s Ctf19^CCAN^:nucleosome complex (PDB ID 6QLD (Yan et al., 2019)) with the same orientation from which the human complex is viewed in panel A. *Right*: In the yeast complex, the 2-fold pseudosymmetry axis relating the LNHDs of CENP-LN is offset relative to the experimentally determined position of the DNA. The LNHD of Chl4^CENP-N^ is displayed with the same orientation as the human’s shown in panel A. **C**) View from PDB ID 6QLD where only the vault and the nucleosome are displayed. **D**) As in panel C, but from a different orientation to emphasize lack of contacts of Iml3^CENP-L^ with the DNA and nucleosome.

**Supplementary Table 1.** Cryo-EM data collection, refinement and validation statistics.

|  | CCAN Krios2 EMDB-14098<br>PDB 7QOO ("dataset I") | CCAN Krios1 EMDB-14099<br>("dataset II") |
| --- | --- | --- |
| **Data collection and processing** |  |  |
| Magnification | 130000x | 105000x |
| Voltage (kV) | 300 | 300 |
| Electron exposure (e-/Å²) | 76.8 | 55.8 |
| Defocus range (µm) | -0.6 to -1.2 | -0.6 to -1.2 |
| Pixel size (native/super-resolution) (Å) | 0.7/0.35 | 0.68/0.34 |
| Symmetry imposed | C1 | C1 |
| Initial particle images (no.) | 140910 | 233598 |
| Final particle images (no.) | 22853 | 44216 |
| Map resolution (Å) | 4.6 | 10 |
| FSC threshold | 0.143 | 0.143 |
| **Refinement** |  |  |
| Initial model used: | AlphaFold,6EQT,4WAU,6WUC,<br>6QLD, 6QLE |  |
| Model resolution (Å) | 3.8 |  |
| FSC threshold | 0.143 |  |
| Model composition |  |  |
| Non-hydrogen atoms | 24952 |  |
| Protein residues | 3086 |  |
| B-factors (Å²) |  |  |
| Protein | 47.8 |  |
| R.m.s. deviations |  |  |
| Bond length (Å) | 0.0035 |  |
| Bond angles (°) | 0.84 |  |
| Validation |  |  |
| MolProbity score | 1.61 |  |
| Clashscore | 5.14 |  |
| Poor rotamers (%) | 1.53 |  |
| Ramachandran plot |  |  |
| Favored (%) | 96.8 |  |
| Allowed (%) | 3.13 |  |
| Disallowed (%) | 0.07 |  |

